# Accelerating protein design by scaling experimental characterization

**DOI:** 10.1101/2025.08.05.668824

**Authors:** Jason Qian, Lukas F. Milles, Basile I. M. Wicky, Robert J. Ragotte, Amir Motmaen, Andrew J. Borst, Rebecca Skotheim, Sebastian Ols, Brian Coventry, Xinting Li, Ryan D. Kibler, Inna Goreshnik, Marc Expòsit, Karin Loré, Lance Stewart, David Baker

## Abstract

Recent advances in de novo protein design have greatly outpaced standard protein biochemistry workflows, and experimental testing has been a bottleneck in the validation of new designs and methodologies. Here, we describe experimental and computational workflows to address the issues of scale, speed and reproducibility of common in vitro protein testing methods, enabling at least an order of magnitude increase in throughput while reducing wetlab time. Semi-Automated Protein Production (SAPP) is a rapid, modular, scalable and cost-effective protocol, enabling up to milligram-scale protein production, and standardized characterization including yield, dispersity, and oligomeric state of hundreds of designs per day, at the cost-equivalent of a few DNA oligos per construct. End-to-end protocol execution takes 48 hours, with about 6 hours spent benchside using mostly standard laboratory equipment. This protocol has become the standard at our institute, providing critical experimental validation for dozens of projects spanning tens of thousands of designs. We showcase the power of the platform by using it to rapidly characterize de novo designed inhibitors of respiratory syncytial virus. Since at least 80% of SAPP’s total cost comes from synthetic DNA, we also developed a scalable demultiplexing protocol (DMX) to leverage oligo pools as input DNA, providing a further 5-fold reduction in costs, enabling >1000 designs to be purified and characterized in arrayed, clonal format at a cost of $5 per construct. By reframing standard molecular biology practices and orchestrating wetlab workflows with partial automation instead of complex end-to-end robotics, these protocols should be widely adoptable, accelerating protein design.

## Main

Recombinant protein expression and purification followed by biochemical characterization is a mainstay of protein engineering. Recent advances in de novo protein design (*1–16*). have greatly outpaced standard protein biochemistry protocols, making experimental validation a bottleneck for protein design efforts. Typical workflows are time, cost, and labor intensive, often requiring multiple weeks for testing a few dozen designs, and lack protocol standardization for consistent data collection, thus precluding large-scale and systematic investigations of the parameter space influencing design success. Pooled assays can test thousands or even millions of designs in parallel (*17–28*), providing a compelling alternative to arrayed testing when large amounts of data are required for a project. However, these assays are usually highly specialized towards measuring one or a few properties (e.g. stability, binding, enzymatic activity), require significant upfront protocol development, and rarely provide information on the manufacturability of individual sequences (expression, solubility, dispersity).

We reasoned that a simple assay-agnostic workflow for the production and initial characterization of hundreds of proteins per day in a standardized and cost-effective manner could significantly accelerate protein design efforts. Moreover, standardizing typical protein characterization data output could enable the use of such experimental data for algorithmic optimization through closed-loop approaches such as active learning (*29–34*). We set out to develop a workflow that generates purified characterized protein starting from gene fragments with greatly reduced execution time, utilizing python scripting to scale and automate key steps in protein purification and characterization using standard laboratory equipment. To reduce the costs further for projects involving characterization of large numbers of designs, we sought to extend the approach to replace gene fragments with much lower cost array synthesized oligonucleotides. We developed workflows for generating purified and characterized protein rapidly from gene fragments (SAPP) and for generating sets of sequence verified fragments from oligonucleotide arrays (DMX) to input into SAPP. These protocols are described in turn below.

## Results

### Semi-Automated Protein Production

The first step in experimentally testing any computer-generated protein is the design of a suitable DNA construct, typically a plasmid. The most economical format for obtaining arrayed synthetic genes encoding new designed proteins is currently DNA fragments, and standard practice has been to clone and sequence verify these constructs. This adds significant cost per design, and is hard to parallelize due to the need for plating individual cloning reactions. We reasoned that the time required could be greatly reduced by omitting this step, and that by standardizing all steps in the purification pipeline could yield both great increases in efficiency and much more globally comparable data (Fig. 1).

**Fig 1.**
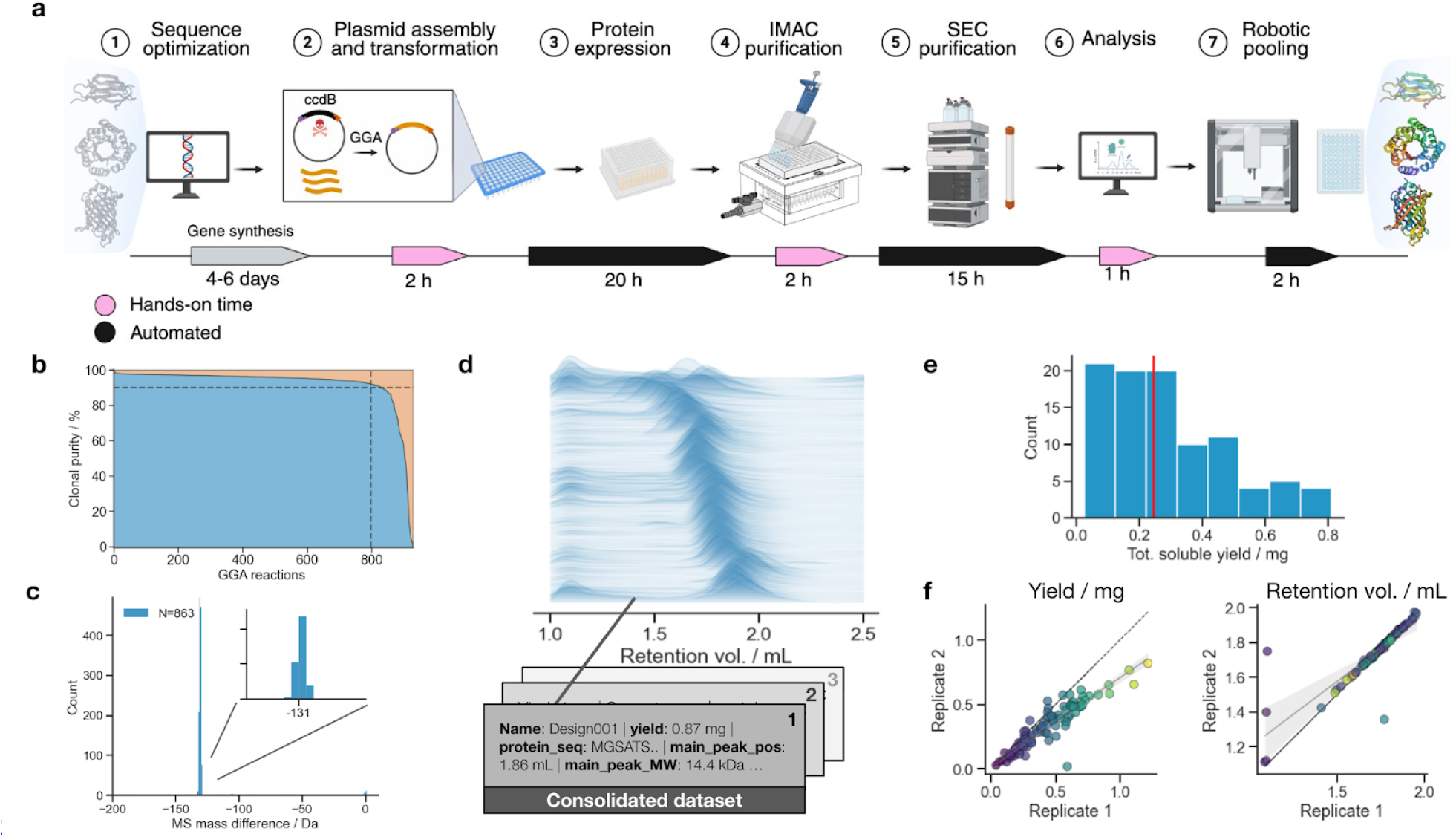
SAPP pipeline and experimental validation. (a) Schematic representation of the SAPP pipeline from cloning to protein expression and purification. (b) NGS data for 929 SAPP samples. Sequencing results of the polyclonal mixtures show that the majority of sequences are correct (blue), with only a small fraction of reactions showing incorrect sequences (orange). Dashed line intersection represents 90% clonal purity. (c) LC-MS data on samples purified by SAPP, showing correct masses, with most proteins lacking their N-terminal Met (-131 Da) as expected in *E. coli* (N=863). (d) Representative results from a SAPP run with 96 samples, showing SEC chromatograms ordered by hierarchical clustering. (e) Histogram showing typical soluble yield in mg for designed proteins purified with SAPP. (f) Parity plots of biological replicates showing total soluble yield in mg (left), and elution of the main peak in mL (right), demonstrating the reproducibility of SAPP.

Plasmid generation involves reverse-translating the amino acid sequence and deciding on expression/purification tags to be included. Often, expression constructs are curated and assembled manually, which is laborious and error-prone. We began by developing software for automated design construction, and an efficient and modular cloning strategy. Starting from a set of desired (designed) amino acid sequences, automated reverse-translation optimizes synthesizability and codon adaptation of the corresponding DNA sequences, and adds standardized adapters for downstream cloning into a suite of compatible vectors for multiplexed cloning; the same design can be subcloned into multiple vectors depending on the desired application(s) (e.g. His-tag, Strep-tag, Avi-tag, NanoBiT, Halo-tag, MBP, FPs, see Supplementary Table 1 for the full list of 25 vectors deposited at Addgene). Large constructs (1.5-3 kb) are automatically split into two compatible fragments for one-pot assembly.

Following construct design, DNA fragments are typically manually assembled into plasmids using a variety of cloning techniques (e.g. Gibson Assembly), followed by transformation into cloning strains, plating, picking of clones, DNA extraction, sequencing, and finally re-transformation of validated plasmids into expression strains for protein production. Sequencing is seen as necessary due to errors resulting from synthesis and cloning. This process adds multiple days and significant costs to any protocol, and scales poorly with colony isolation being a bottleneck. We reasoned that by optimizing a standardized cloning approach we could achieve efficiencies sufficient to bypass these steps. Design protein sequences are batch reverse-translated using custom software, optimizing user-defined codon adaptation index and suppressing alternative start sites while ensuring synthesizability by DNA manufacturers. This process appends cloning adapters for compatibility with a range of entry vectors, and automatically splits large genes into two fragments for one-pot assembly. DNA constructs are ordered as linear gene fragments from commercial manufacturers (list price $0.07/bp) and cloned into the desired receiving vector(s) by Golden Gate Assembly (GGA) (*35*, *36*) followed by direct transformation into an *E. coli* expression strain. Target vectors contain a lethal *ccdB* gene (*37*) that is excised and replaced by the design during GGA, ensuring background suppression and thus circumventing the need for colony isolation. The whole process from linear DNA to inoculated cultures is optimized to take 2 hours for up to 192 reactions. While polyclonality due to synthesis or cloning errors is a risk with this approach, sequencing analysis of 929 GGA reactions with insert size below 314 bp showed that 89.3% had a clonal purity ≥ 90% (Fig. 1b). Since protein expression can further bias polyclonal mixtures due to fitness effects resulting from variants, we performed mass spectrometry analysis of 863 purified proteins to analyze the extent of any possible contamination. This revealed the expected mass within ± 1 Da for 90% of all products (777 / 863) (Fig. 1c). These results indicate that our optimized molecular biology workflow is sufficiently reliable to ensure that the correct clone is the dominant construct in the expression pool in most cases, thus circumventing traditional multi-day cloning protocols. While highly efficient, our protocol may still occasionally yield false negative results, and thus is primarily intended for screening purposes.

When chromatographic protein purification and sizing is intended, shake-flask cultures are the norm. However this approach is difficult to scale beyond a few dozen. Moreover, protein expression is often induced chemically (e.g. IPTG) when cultures reach a certain density, which is inherently challenging to parallelize. We explored the ability of auto-induction media formulation to yield sufficient protein from microplate cultures to streamline multiplexing and reduce human intervention. Cultures are grown in 96-well deepwell plates in autoinduction media at constant temperature, avoiding the need for any intervention between inoculation and harvest (*38*).

When screens are performed in microplate formats, the end-point is usually an SDS-PAGE gel to assess expression, and this information is then used for selecting candidates for scale-up and full chromatographic characterization. While gel quantification is possible, it does not inform on the native properties of proteins (oligomeric state, dispersity), and this two-step approach is also more time-consuming. We reasoned that by optimizing protein purification we could efficiently perform SEC in a preparative fashion on *every* sample, combining characterization and purification into a single process. Moreover, SEC chromatograms are inherently more data-rich, generating more useful information about both failed and successful designs. Following chemical lysis, proteins are purified from their soluble fractions (a protocol for purification from inclusion bodies is included in Supplementary Information), first by affinity chromatography (typically Ni-NTA) in 96-well format, followed directly by size-exclusion chromatography (SEC) on a liquid chromatography system equipped with an autosampler. The process is optimized to take 2 hours from cell harvest to starting SEC, which is run overnight (15 hours for 192 samples, <5 minutes per sample, see Methods).

Chromatographic results are typically analysed in vendors’ proprietary software using GUI and manual assessments of individual traces. This process scales poorly beyond a few dozen, and is not systematic. We developed open-source software for analyzing thousands of elution profiles. Chromatograms are automatically analyzed, and the experimental profile for each design (yield, polydispersity, estimated molecular weight, see Supplementary Information) is outputted to a single open format file (Fig. 1d).

Beyond chromatogram analysis, fraction picking for pooling, concentration measurement and normalization are also time-intensive manual tasks. Here we integrate an affordable and programmable liquid handling robot into our analysis code to automate the task. The analysis pipeline automatically identifies peaks for pooling and estimates concentrations by integrating A280 absorbance of SEC chromatograms based on sequence-specific extinction coefficients. This information is then used to automatically generate the code necessary to program a low-cost open-source liquid handling robot for the task of simultaneously pooling fractions and normalizing concentrations.

Outcomes of SAPP are consolidated 96-well plates containing the purified proteins (typical yields; 0.1–0.5 mg at 10–100 μM, Fig. 1e), and a single dataframe containing all the experimental data consolidated in one file. By focusing on an optimized molecular biology workflow and not over-relying on automation, SAPP remains modular, flexible and accessible; a user may elect to process a subset of a 96-well plate, or multiple plates with only minimal additional time (the main bottleneck being SEC, with a capacity of ∼300 samples per chromatographic instrument per day). Keeping all steps in a plate format also eliminates container tracking overhead, and reduces single-use plastic waste. Replicate experiments demonstrate that SAPP is robust (Fig. 1f), enabling quantitative analyses on the data generated by the protocol for machine learning applications.

As a demonstration of the protocol for protein design/engineering efforts we applied SAPP to characterize ProteinMPNN-redesigned sequences of fluorescent proteins (FPs), with the goal to rapidly find more stable FPs (Fig. 2). Crystal structures and AlphaFold2 (*39*) predictions from different FPs were selected as starting points. During sequence design, residues in contact with the chromophore, and residues that are highly conserved (top 50-70% conserved in MSA) were kept fixed, as this was shown to be important when applying ProteinMPNN to the redesign of natural protein (*40*). Overall, sequence identities of designs to WT ranged from 50 to 80% with a Levenshtein distance of up to 50 compared to their respective WT (Fig. 2b). We selected 96 constructs for testing, and obtained all experimental results within a week, identifying many designs that have comparable or better yield than the WT (Supplementary Fig. 1). Some exhibit higher thermostability than their WT counterpart (e.g. muGFP), some re-designed sequences remained monomeric after boiling (95° C, 1h) as assessed by SEC (Fig. 2d,e; Supplementary Fig. 2), and also retained absorbance, indicating that both the chromophore and protein remained intact, while also showing altered spectra. For some SYFP2 variants, absorbance spectra changed towards a longer Stokes shift than the WT, showing that surface redesign influences the photophysical properties of these fluorescent proteins (Fig. 2d,e). In conclusion, using SAPP enabled 96 FP variants with large edit distances to be characterized within a week, providing detailed biochemical information (yield, dispersity, oligomeric state, absorbance and fluorescence) for each design.

**Fig 2.**
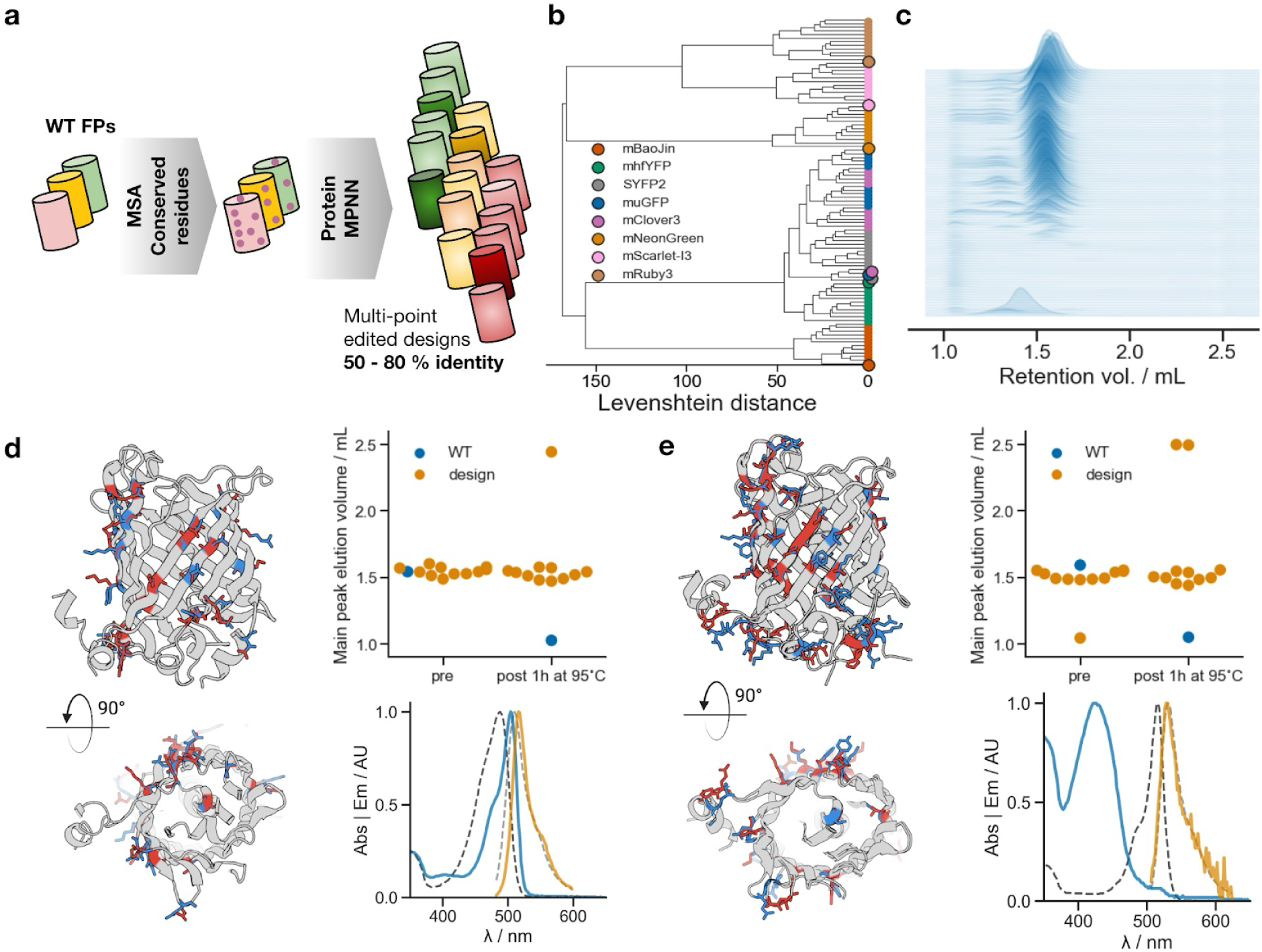
SAPP for Fluorescent Protein Sequence Redesign with ProteinMPNN. (a) Non-conserved surface residues of fluorescent proteins were redesigned using ProteinMPNN. (b) A dendrogram, based on Levenshtein distance, displays 96 characterized sequences from SAPP, color-coded by FP template. Wild-type sequences are marked with circles. (c) Aggregate SEC traces of the 96 designs reveal distinct elution peaks near the anticipated molecular weight for over half of the variants. (d, e) For designs 4389 and 4468, the redesigned amino acid positions (red) are highlighted on the structures of their wild-type templates, muGFP (d) and SYFP2 (e), with the original amino acids shown in blue. The top plots show elution profiles post-purification and after a 1-hour incubation at 95°C for all variants of that parental sequence. The bottom plots present the absorbance and emission spectra of the designed variants shown on the left (colored lines) and their corresponding wild-types (dashed lines), indicating changes in photophysical properties.

### RSV binders

Screening binding proteins in diverse oligomeric states has important applications within cell signalling and pathogen neutralization (Fig. 3a). In cell signalling, lower-order oligomers (dimers, trimers) can be used to induce receptor dimerization/trimerization to agonise the target pathway (*41*, *42*) while high valency oligomers can induce signalling in pathways that require higher order cross linking, such as T cell receptor 4-1BB (*43*). For virus neutralization, surface exposed viral glycoproteins are amenable to symmetry matching (*44*, *45*), while the regular spacing and repetitive structure of non-enveloped viruses and some enveloped viruses (e.g. alphaviruses) could be targeted with high valency oligomers (Fig. 3a). Unfortunately, the non-covalent nature of oligomers makes multimeric constructs inherently incompatible with display technologies such as yeast surface display (YSD). We reasoned that we could apply the SAPP pipeline to rapidly screen 116 oligomeric constructs of de novo designed minibinders to identify correctly-assembled oligomers, and then screen for the optimal geometry in direct functional assays. This would enable the rapid identification of optimized oligomeric constructs without needing to design bespoke oligomers for each new application while still exploring diverse valencies and spacing between binding domains towards improved inhibitory potential.

**Fig 3.**
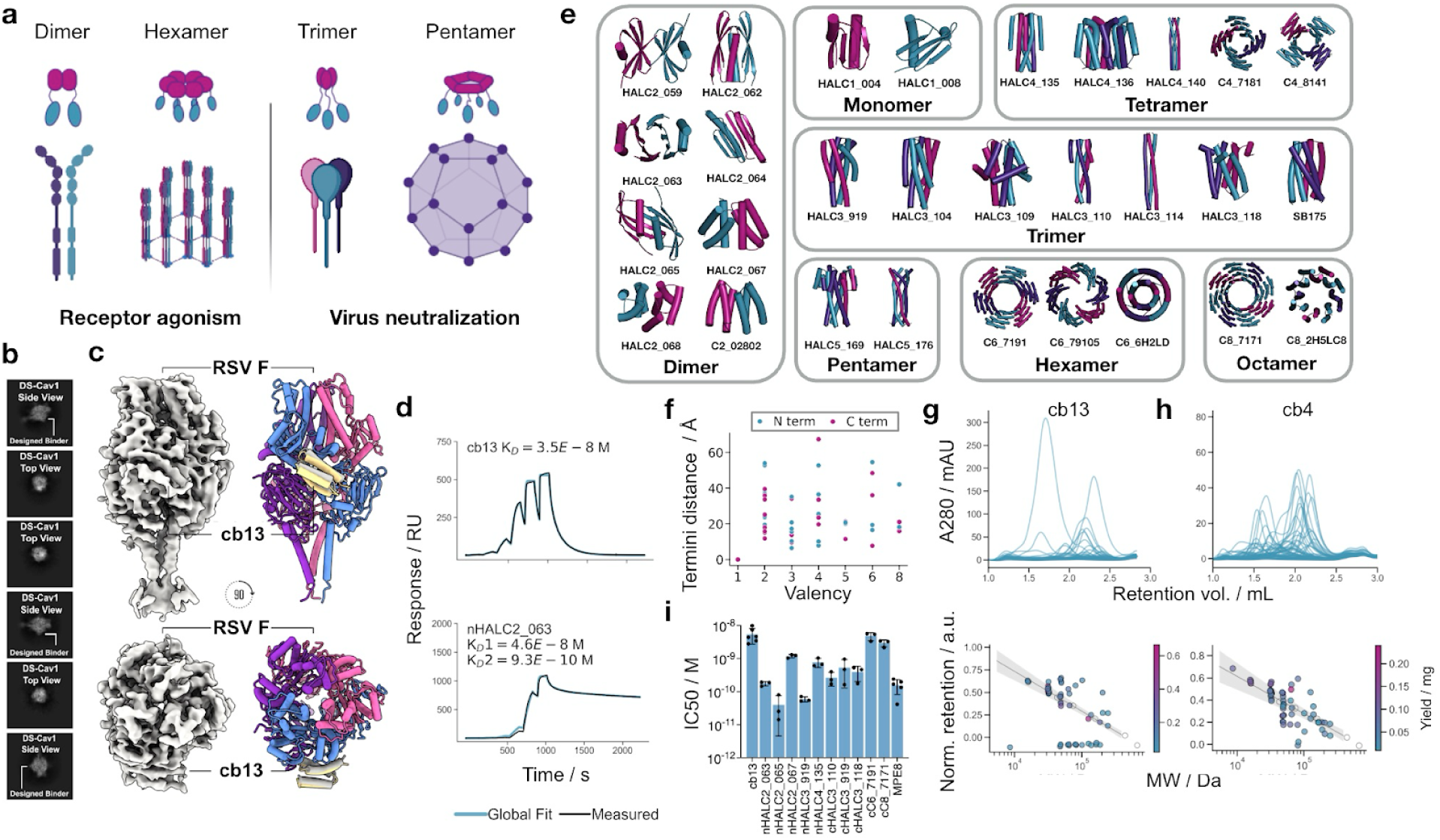
RSV oligomer screening. (a) Lower-order oligomers (dimers, trimers and tetramers) can be used to create agonists for receptor signalling by bringing two receptor chains together or to make symmetry matched neutralizing molecules that bind to typical trimeric and tetrameric viral glycoproteins. Higher order oligomers can be used to crosslink larger networks of surface receptors to induce signalling or to target symmetry on the viral capsid. (b) 2-D classification of cb13 bound to SC-DM, a sequence variant of RSV F glycoprotein. (c) 4.63 Å Cryo-EM density map and model of cb13 bound to RSV F (yellow), as compared to the computational design model (grey) (Cα RMSD = 1.9 Å). (d) Surface plasmon resonance measurements of monomeric or multivalent cb13 constructs binding the RSV F immobilized on a CM5 chip through amine conjugation across a 6 step concentration series beginning at 320 pM and increasing 5-fold with each subsequent injection. 1:1 binding was used to fit the cb13 binding curve while the heterotypic ligand was used for nHALC2_063 where the oligomer may engage one or more RSV F copy at a time. (e) Overview of the oligomeric constructs included in the screening library ranging from dimers to octamers alongside two monomer controls. (f) Geometric diversity of the screening library assessed by valency and distance between the termini of the oligomers. Pink and cyan indicate the spacing between the oligomer N and C termini in each chain, respectively (g-h) SEC trace overlay of cb13 (g) and cb4 (h) fused to each of the oligomers within the screening library (upper). Elution volume relative to a molecular weight standard to determine if the oligomers have assembled into their intended multivalent structure (lower). The shaded area indicates the 99% confidence interval for elution volume based on the molecular weight standards. (i) 50% inhibitory concentration (IC50) for cb13 monomer and oligomeric constructs in live RSV virus in neutralization assays. Prefixes “n” and “c” in front of design names mean N- and C-terminal binding domain, respectively. Points indicate IC50s from individual experiments and error bars indicate standard deviation.

We used this approach to explore oligomerizing a designed binder of respiratory syncytial virus (RSV) F protein to improve neutralization. We used the Rosetta-based rotameric interaction field (RIF) method to generate binders to site III of the F protein. The top ∼31,000 designs were obtained as chip-based synthetic DNA, screened through yeast surface display, and optimized through site saturation mutagenesis. The best resulting design, cb13 (Fig. 3b,c), had an affinity of 35 nM for RSV F protein (Fig. 3d). A cryo-electron microscopy structure of cb13 bound to RSV F confirms that the molecule binds as designed, with 1.9 Å RMSD across all Cα of the binder when aligned on the target (Fig. 3b,c, Supplemental Fig. 7, Supplementary Table 9).

We built a 54-member oligomer library (plus 4 monomer controls for 58 total constructs) using a selection of 27 oligomers, each with either an N- or C-terminal fusion, made through AF2-based hallucination (*5*) and Rosetta (*42*, *46*). These spanned valencies from C2 - C8 with binding domain spacing of 10Å - 65Å (Fig. 3e,f). With 58 entry vectors arrayed on an Echo acoustic liquid handler-compatible plate, the SAPP pipeline was then used to clone, express and purify oligomer variants of cb13. The pipeline rapidly identified well-expressing and assembled oligomers for prioritization (Fig. 3g). Because the oligomer vectors use the same overhangs as the other SAPP expression vectors, the same gene fragment can be used for initial hit identification and downstream oligomer screening.

We identified 19 cb13-oligomer fusion constructs with moderate to high expression (>0.1 mg per 4 mL of E. coli culture) that assembled to their expected oligomeric state (Fig. 3g), as defined by an elution volume within the 99% confidence interval of expected elution volume on an S200 5-150 column. We then tested monomeric cb13 and a subset of these oligomeric cb13 constructs in virus neutralization assays. The oligomeric constructs vastly outperform monomeric cb13 with nHALC2_065 and nHALC3_919 having IC50s of 40 and 59 pM respectively, compared to 5.4 nM for their monomeric counterpart (Fig. 3i). These oligomers had IC50s similar to, or better than, the potent benchmark monoclonal antibody MPE8 (*47*), which targets the same epitope of RSV F and had an IC50 of 156 pM (Fig. 3i). We confirmed that the improved neutralization is due to slowed dissociation as a result of avidity effects by comparing apparent binding affinities of the monomeric cb13 to nHALC2_063 where we see a slowing of dissociation from RSV F in the multimeric form (Fig. 3d). These data also highlight the utility of screening diverse oligomerization geometries. In neutralization assays, nHALC2_065 is ∼25-fold more potent than an alternative dimer, nHALC2_067 (IC50s: 40 pM vs 1 nM), despite having identical binding domains (cb13) and valency (dimers). Likewise, the same trimerization domain, HALC3_919, is 10-fold more potent with the binding domain at the N terminus (59 pM) than the C terminus (530 pM). Higher order oligomers (C6 and C8) do not show meaningful improvements over the cb13 monomer (Fig. 3i).

We also oligomerized cb4, a sequence variant of cb13 containing seven point mutations with marginally weaker neutralization. The success rate was higher than for cb13 (33 of 58 constructs assembled to their target oligomeric state) showing that oligomerization success can be dependent on the binding domain in the fusion constructs: not all binding/oligomerization domains are mutually compatible (Fig. 3h). Applying SAPP to oligomer screening identifies compatible pairs of oligomerization and binding domains and enables rapid generation of diverse oligomers to probe how geometry shapes function.

### DMX

The highly optimized SAPP protocol has shifted the bottleneck of large-scale testing from protein expression/purification to the cost of DNA synthesis (Supplementary Fig. 3, Supplementary Table 2). While oligo pools provide an economical alternative to gene fragments, they are not immediately suitable for the vast majority of downstream biochemistry workflows, which are often performed in arrayed format. To take advantage of the low cost of oligo pools while retaining the modularity of SAPP, we developed a protocol to convert DNA oligo pools into clonal genes of interest (GOI) using a process that is cost-effective, scalable, and generates sequence-verified constructs. During implementation we focused on: (i) a DNA barcoding strategy that can easily be scaled to thousand of variants; (ii) a demultiplexing strategy that is agnostic to GOI length; (iii) total end-to-end costs (for DNA synthesis, barcoding reagents and sequencing) considerably lower than the cost of individual gene fragments; (iv) seamless integration with SAPP for downstream characterization. Protocols for the demultiplexing of pooled variants using barcoding and sequencing have been implemented previously (*48–52*) but none explicitly addressed all these requirements. The DMX pipeline is a 5-day protocol that leverages an isothermal DNA barcoding strategy and long-read sequencing to produce thousands of sequence-verified clonal cultures ready for processing with SAPP or any other testing modalities that require arrayed DNA (Fig. 4a).

**Fig 4.**
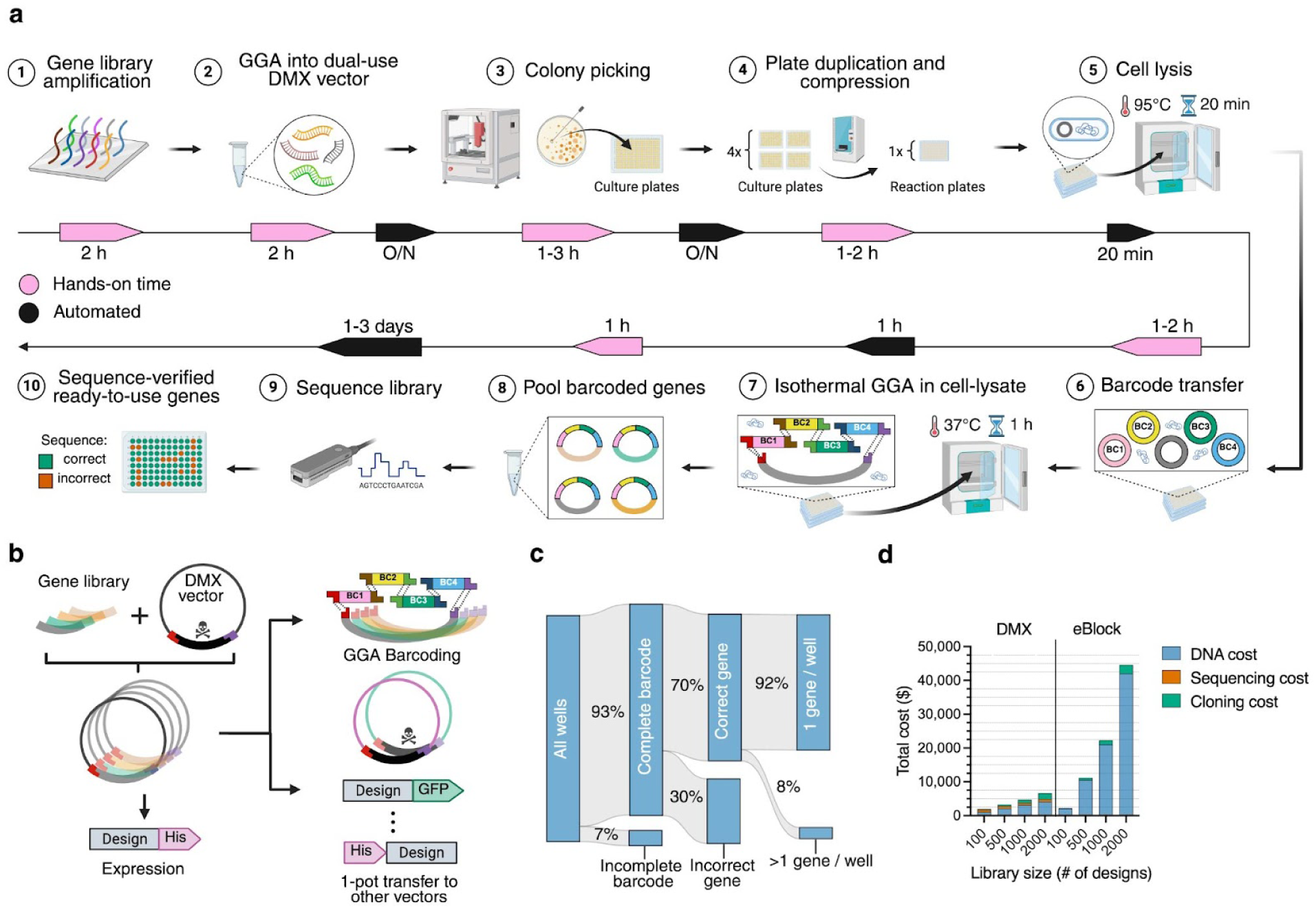
DMX pipeline. (a) Schematic representation of the DMX pipeline from oligo pool library to sequence-verified ready-to-use clonal bacterial cultures. (1) DNA library is amplified from the chip; (2) library is cloned into DMX vector using GGA, transformed and plated; (3) colonies are picked into culture plates; (4) culture plates are duplicated and compressed 4-to-1 into reaction plates using ECHO; (5) cultures in reaction plates are heat-lysed at 98 °C for 20 minutes; (6) barcodes are transferred into each well of the reaction plates; (7) GGA reagents are transferred and barcodes and GOI are assembled by incubating reaction plates at 37 °C for 1 hour; (8) pool all GGA products; (9) sequence barcoded library using ONT; (10) re-array sequence-verified clonal variants into new plates. Pink and black arrow boxes represent hands-on time and automated steps, respectively. (b) Schematic representation of multi-use vector cloning for pool entry (BsmBI), barcoding (BsaI), and GOI excision (BsaI) if desired for further subcloning into other vectors. Skull represents the *ccdB* gene. (c) Sankey diagram showing the efficiency of the DMX pipeline (N=4,608 wells). (d) Cost comparison between sourcing GOIs using DMX from chip or arrayed DNA fragments, broken down into DNA synthesis, sequencing, and cloning expenses.

When sequencing variants from oligo pools, variant validation and downstream characterization are often decoupled, which adds time and overhead. Instead we aimed to use a dual purpose pool entry vector, and transformation directly into an expression strain to couple barcoding/sequencing with strain generation. The DMX entry vector was engineered with triple GGA capabilities, to allow both pool entry, barcoding, and optionally GOI excision for further subcloning in an almost scarless manner if desired (Fig. 4b, Supplementary Fig. 4). After amplification from a pool, the DNA library is cloned into the DMX vector by GGA, transformed into expression-ready competent *E. coli* cells by electroporation, and plated on LB-agar for colony generation.

Following isolation into 384-well ECHO plates using a colony picker, GOIs are barcoded. This step is typically achieved by dual-indexed PCR, which limits throughput, and requires large numbers of thermocyclers when processing thousands of samples. To overcome this limitation, we developed an isothermal barcoding strategy based on GGA that ligates barcodes with the GOI directly in bacterial lysate. Our approach uses four sets of position-specific barcodes comprising 24 unique molecular identifiers (UMIs) each, allowing for >330k wells to be tagged from a single 96-well plate of UMIs (24^4) (Supplementary Table 3,4). This expansive barcode capacity exceeds the dual-barcoding approach typically used in individual plates (*48*, *51*) or the positional barcodes used in the bacterial mating system (*52*). Importantly, the barcoding reaction is isothermal, obviating the need for thermocyclers, and our plasmid-derived barcodes can be cheaply produced, making the process simple and scalable.

Once barcoded, samples are pooled, GOIs are mapped to their barcodes using next-generation sequencing. This is typically done with Illumina short-reads, which limits the sequencing length of GOIs (e.g. 300–600 bp amplicon lengths). We leverage nanopore long-read sequencing to overcome this limitation, making barcode mapping length-agnostic, and enabling pools with longer genes to be demultiplexed, such as those obtained by Dropsynth (*53*), paired assembly (*54*, *55*), or multiplexed GGA of long GOIs (*56–58*).

An automated bioinformatics pipeline performs quality control on the sequencing results; basecalling the reads from the nanopore sequencer, filtering based on quality, barcode demultiplexing, aligning the demultiplexed reads to their references, and finally generating a consensus sequence (see Supplementary Methods). This returns a non-redundant pick-list of wells, which are re-arrayed by the ECHO in a final step. Outcomes of DMX are arrayed cultures of clonal sequence-verified plasmids in expression-competent *E. coli* cells ready for downstream processing with the SAPP pipeline.

To demonstrate the cost-effectiveness and scalability of the DMX pipeline, we obtained an oligo pool encoding a library of 1,500 de novo designed protein with lengths ranging from 275 to 400 bp. We successfully recovered 78% of the library variants as clonal, sequence-verified constructs by screening 4,608 clones (3x the library complexity), approaching the theoretical maximum recovery rate of 92% for the given library complexity, sampling depth, insert length, synthesis error and synthesis bias reported by Twist (Fig. 4c and Supplementary Fig. 5).

We developed the DMX protocol using automated instruments such as the ECHO liquid handler and colony picker, but it can also be implemented without access to such equipment using low-cost robotics (*59*) like an OT-2 or manual picking and multichannel pipettes. For large demultiplexing campaigns, the optimal library size is about 2,000 genes, requiring the sampling of about 8,000 colonies (20 x 384 well plates) to achieve >95% recovery (Supplementary Fig. 5). For smaller campaigns, libraries of 500 genes can be demultiplexed by manually picking 1500 colonies in order to maintain the same recovery rate. This approach is five and eight times more cost-effective (including reagent and sequencing costs) than ordering 500 and 2000 individual gene fragments, respectively (Fig. 4d, Supplementary Table 5), and ensures clonality of tested variants, while only adding five days to the pipeline. Thus it is ideally suited for large design campaigns where the time overhead is absorbed over many constructs, while the polyclonal approach from SAPP is better suited for smaller campaigns (∼100-200 designs) where faster turnaround time is desired.

## Discussion

The modular SAPP workflow dramatically accelerates the experimental characterization of designed proteins. SAPP combines throughput-optimized molecular biology steps, an automated analysis pipeline, and optionally open-source robotics to purify and characterize hundreds of designs per day in array format with only a few hours spent at the bench, with typical yields being sufficient for most downstream biochemical assays (e.g. binding, enzymatic activity, thermostability, cell assays, electron microscopy). Complete end-to-end execution takes 48 hours (excluding commercial DNA synthesis), enabling rapid iteration cycles at a total cost per design equivalent to a few DNA oligos (for a detailed calculation see Supplementary Information). Two key aspects enable the speed and efficiency of SAPP. First, the use of modular cloning and a background-suppressing cassette makes direct-to-expression from commercial gene fragments sufficiently reliable for screening applications, bypassing the standard procedure of clone isolation and sequencing. Second, streamlining the SEC workflow allows for data-rich information generation on every design, which can be automatically analysed and consolidated into an open format for quantitative analyses between design campaigns. By optimizing common experimental techniques over complex robotics, this protocol can be adopted by most labs.

We applied this framework to the rapid development of potent multimeric de novo RSV F inhibitors, which required screening of 116 constructs. This yielded designs that are on par with a highly potent RSV site III mAb, MPE8. The speed of the SAPP protocol reduces the time to lead candidate identification, and enables the exploration of geometry and valency as an added dimension during inhibitor optimization.

To further reduce the cost of per-design testing, we developed the DMX pipeline to convert DNA oligo pools into arrayed cultures of sequence-verified plasmids. The scalability of DMX is primarily driven by a combinatorial in-lysate isothermal barcoding framework, which eliminates the need for thermocyclers, and by nanopore long-read sequencing, which offers a low financial entry point and is compatible with applications involving longer GOIs. The protocol achieves high recovery rates of sequence-verified constructs, and integrates seamlessly with downstream workflows like SAPP.

These protocols have already been used to experimentally produce and characterize tens of thousands of de novo designed proteins at the Institute for Protein Design, ranging from small mini-protein binders against cell surface receptor of therapeutic interest, de novo enzymes, and symmetric protein assemblies hundreds of kDa in size (*1*, *4*, *5*, *8*, *10*, *40*, *43*, *60–72*). These workflows both democratize and accelerate protein design, making large-scale experimental testing more affordable and accessible. We envision that the integration of scale, standardization, and speed will contribute to the development of new protein design models informed by experimental data, as well as enabling active learning approaches in protein design. Collectively, SAPP and DMX present an experimental counterpart matched to the rapid pace and scale of recent advances in computational protein design.

## Methods

The list of equipment, items and components necessary to the execution of the protocol (including suppliers and catalog numbers) can be found in Supplementary Table 6 and 8 for SAPP and DMX respectively. The composition of SAPP buffers can be found in Supplementary Table 7. Chemicals were purchased from Millipore-Sigma or Fisher Scientific unless stated otherwise. DNA fragments were purchased as eBlocks from Integrated DNA Technologies (IDT). Oligo pools were purchased from Twist Bioscience.

### DNA fragment generation

Protein sequences are first reverse-translated to DNA sequences using a custom python script that takes a FASTA file or a list of PDBs as input. Codon optimization is performed with DNA Chisel (*73*) using a mixed objective function that optimizes user-defined codon adaptation index (CAI), minimizes repeat sequences, avoids BsaI restriction endonuclease recognition sites, and suppresses alternative start sites. The script also ensures synthesizability on the fly by using IDT’s SciTools Plus API, and if the original sequence cannot be made, the objective function is gradually modified by up-weighting the repeat sequence cost parameter until synthesizability is achieved. If the reverse-translated sequence is longer than a user-specified length (default: 1500 bp, IDT’s eBlock maximum), the sequence is automatically split into two fragments for one-pot Golden Gate Assembly (GGA) (*35*). The 4 bp fragment overhang is chosen from a predefined list that maximizes orthogonality to the entry vector GGA sites (5’->3’: AGGA and TTCC) in order to ensure assembly fidelity (*74*). Finally, overhangs for GGA containing BsaI recognition sites are appended to the 3’ and 5’ ends, and random filler sequences added to the outer parts if the sequence is less than 300 bp (IDT’s eBlock length minimum). The script outputs a formatted spreadsheet that is directly uploaded to IDT for ordering as eBlocks ($0.07/bp, 1-3 business days turnaround), and an ECHO transfer protocol for automated cloning (see below). DNA fragments are ordered resuspended in IDTE buffer (10 mM Tris, 0.1 mM EDTA, pH 8.0) in either 96-well PCR plates (for manual cloning), or ECHO-qualified 384-well plates (for robotic-assisted cloning). The whole process is fully automated, and going from a list of amino acid sequences or PDBs to placing the gene fragment order only takes minutes.

### Cloning

Entry vectors are propagated and produced in bulk from *ccdB*-resistant NEB Stable *E. coli* chemically competent cells (NEB). Following transformation according to the manufacturer’s protocol (see Supplementary Information), the transformants are grown overnight at 37 °C on LB-Agar containing 50 μg/mL of kanamycin sulfate. Single colonies are used to inoculate 50 mL of liquid LB cultures (in 250 mL baffled conical flasks) containing 50 μg/mL of kanamycin sulfate, and grown overnight at 37 °C with shaking at 250 rpm. Plasmids are purified using a ZymoPURE II Plasmid Midiprep Kit (Zymo Research) according to the manufacturer’s protocol, and verified by full plasmid sequencing (Plasmidsaurus). Typical yields are 50–100 μg, which is enough for about 2,000–4,000 GGA reactions.

GGA reactions are assembled with an ECHO 525 acoustic liquid handling robot (Beckman Coulter) by using the transfer protocol outputted during DNA fragment generation. A master mix containing BsaI-HFv2 (NEB), T4 DNA ligase (NEB), T4 DNA ligase buffer (NEB) and the entry vector is prepared first, and the ECHO is used to combine master mix and DNA fragments into 96-well PCR plates (1 μL final volumes). Reactions are incubated at 37 °C for 15 minutes, followed by 5 minutes at 60 °C. GGA products are directly transformed into an *E. coli* expression strain by adding 6 μL of BL21(DE3) chemically competent cells (NEB) to the 1 μL reactions. Transformations are performed in 96-well PCR plates (see Supplementary Information), and the transformed cells are used to inoculate expression cultures directly (transformation efficiency = 858 ± 443 colony forming units (CFU) (mean ± standard deviation, *N* = 12)). Going from linear DNA to inoculated cultures takes about 2 hours for 192 GGA reactions, and only requires an extra 15 minutes for every additional 96-well plate.

If an ECHO robot is not available, we recommend performing 5 μL cloning reactions; first by aliquoting 4 μL of master mix into each well, followed by the addition of 1 μL of eBlock using a multichannel pipette (5 μL total volume). After GGA, 1 μL is used to transform 6 μL of cells as described above.

### Protein expression

Protein expression is performed in round bottom 96 deepwell plates filled with 1 mL of simplified auto-induction media (*38*) (TB II, MP Biomedicals, supplemented with glycerol [5 g/L], glucose [0.5 g/L], lactose [2 g/L], MgSO_4_ [2 mM], and kanamycin sulfate [50 μg/mL]). Transformation reactions are split 4-fold and used to inoculate 4 x 1 mL of expression cultures directly, followed by incubation at 37 °C for *at least* 20 hours at 1,000 rpm on a 2.5 mm orbital shaker (NB smaller orbital paths appear to negatively impact expression, most likely due to reduced aeration (*75*), which likely impacts the glucose to lactose metabolic switch necessary for auto-induction).

### Affinity chromatography

Cells are harvested in 96 deepwell plates by centrifugation (4,000 x g, 5 minutes), the media discarded, and cells resuspended in lysis buffer (B-PER, Thermo Fisher Scientific, supplemented with lysozyme [0.1 mg/mL], PMSF [1 mM], Benzonase [25 U/mL], 100 μL per 1 mL of culture-equivalent pellet). Lysis is left to proceed for 15 minutes at 37 °C under agitation (1,000 rpm) before pooling the lysates (4 x 100 μL) into one 96 deepwell plate and clearing debris by centrifugation (4,000 x g, 15 minutes). Proteins are purified from the soluble fraction by binding to Ni-NTA resin (50 μL of resin bed per well, added to 96-well fritted plates (25 µm PE frit, Agilent Technologies)), followed by three wash cycles using a plate vacuum manifold (3 x 400 μL, 20 mM Tris, 300 mM NaCl, 25 mM imidazole, pH 8.0). Proteins are eluted from the resin by addition of 200 μL of elution buffer (20 mM Tris, 300 mM NaCl, 500 mM imidazole, pH 8.0), and sterile-filtered by centrifugation into a 96-well filter plate (0.22 μm pore-size) mounted on a receiver plate. The whole process process is optimized to take under 2 hours from cell harvest to injection-ready eluates for up to two 96-well plates, with only minimal extra time necessary for additional plates.

### Size-exclusion chromatography

Filter-sterilized eluates are purified/analyzed by size-exclusion chromatography (SEC) into PBS on either a Superdex 75 (70 kDa MWCO) or Superdex 200 (600 kDa MWCO) column (5/150 with 3 mL column bed volume, Cytvia) by injecting 100 μL at a flow-rate of 0.65 mL / minutes (<5 minutes per sample, ∼15 hours for 192 samples). Chromatography is performed either on a Agilent 1260 Infinity II Bio-inert system equipped with multisampler and fraction collector (fractionation into 384-well plates), or a Cytvia ӒKTA Pure system equipped with an autosampler and using a modified flow-path (fractionation into 96-well plates, see Supplementary Information for details). Chromatograms are batch-analyzed by a custom python script (see below) which also generates code to program an OT-2 liquid handling robot (Opentrons) for cherry-picking, pooling and normalizing fractions. The end-point of the protocol is a set of 96-well plates containing SEC-purified and concentration-normalized proteins (typically a few hundreds microliters at single to double-digit micromolar concentrations).

### Data processing

The integrated data-analysis pipeline starts by generating plasmid maps from the ordered DNA fragment sequences and specified entry vectors. The ORF is used to generate the expression sequence (designed sequence plus vector-specific tags), and calculate protein parameters (molecular weight, molar extinction coefficient, charge, isoelectric point, etc). Batch-exported chromatograms are analyzed using NumPy (*76*), SciPy (*77*), and Pandas (*78*, *79*), to detect peaks in the protein absorbance signal at 280 nm and convert their retention volume to approximate molecular weight compared to calibration curve assembled from standards (LMW [S75], or HMW [S200], Cytvia). In addition, by integrating the absorbance signal to determine protein concentration in each fraction, the following information can be extracted for each design: total soluble yield, polydispersity, estimated molecular weight and aggregation state of each peak, desired fractions to pool. The data is exported into a standardized open format dataframe (HDF5 (*80*)).

### Mass spectrometry

Intact mass spectra were obtained by reverse-phase LC/MS on an Agilent G6230B TOF using an AdvanceBio RP-Desalting column with a fast gradient of 90:10 (A:B) to 5:95 (A:B) over 2 minutes (A: H2O with 0.1% Formic Acid, B: Acetonitrile with 0.1% Formic Acid), and subsequently deconvoluted with Bioconfirm using a total entropy algorithm.

### Next-generation sequencing

A retrospective analysis of designs ordered as DNA fragments (eBlocks, IDT), assembled, cloned and transformed directly as described above was performed to assess clonal purity (Fig. 1b). Stabs from 929 glycerol stocks (made from LB out-growth cultures with antibiotics) were pooled, and the inserts amplified by PCR using Q5 (NEB), followed by purification using AMPure XP magnetic beads (Beckman Coulter) and analyzed on a 4200 TapeStation (Agilent). Paired-end sequencing was performed using NextSeq2000 (Illumina), basecalled and filtered for quality. Reads R1 and R2 were extracted using SAMtools (*81*), followed by pairing using PEAR-0.9.11 with options-j 20-p 1.0 (*82*). Index sequences of 929 reference sequences were built and aligned to the PEAR output using Bowtie2 (*83*), generating 31 million aligned reads, which were then converted to BAM, sorted and indexed using SAMtools. From the CIGAR strings, clonal purity scores were calculated by dividing the number of perfect matches by the total number of filtered reads for each reference sequence using a custom python script.

### Fluorescent protein sequence redesign

Fluorescent protein (FP) sequences were generated using the protocol reported by Sumida et al. (*14*) In brief, WT FP sequences were extracted from FPbase (*84*), and their structures predicted with AlphaFold2 using MSAs generated from UniRef30_2022_02 using HHblits (*85*). Per position conservation was assessed from the MSAs, and either 50% or 70% of the most preserved residues were kept fixed during ProteinMPNN sequence redesign. Additionally all chromophore forming residues, and all core-facing residues (as assessed by a SASA < 2.0 Å^2 using pyrosetta (*86*, *87*)) were also fixed. ProteinMPNN was run at T=0.01, 0.1, 0.2, 0.3, 0.5, and designed sequences predicted with AlphaFold2 using a shallow MSA of up to 15 sequences. Designs were selected based on the following metrics: AF2_plddt_mean > 0.8, AF2_ptm > 0.80, AF2_pae_mean < 12.0, AF2_tm_score to WT > 0.7, AF2_tm_rmsd to WT < 1.5, AF2_sap_per_res < 0.4. Using TM-align (*88*), SAP score (*89*), and AF2 confidence metrics.

### DMX barcode design

Each of the four position-specific barcode sets comprise 24 UMIs. These 25-bp UMI sequences were taken from (*90*), and further filtered for: pairwise Hamming distance >5, absence of homo-polymers >4 bp, and absence of common restriction enzyme sites (AgeI, BamHI, BbsI, BsaI, BsmBI, BtgZI, EcoRI, HindIII, KpnI, NheI, NotI, PacI, PaqCI, PstI, PvuI, SacI, SalI, SapI, SmaI, XbaI, XhoI). Primer sequences were then appended to each UMI (balanced G/C % for 62 °C Tm), GGA overhangs and BsaI cut sites (Supplementary Table 4). Barcodes were ordered subcloned from IDT as MiniGene 25-500 bp inserted into pUCIDT (Amp) Golden Gate vector.

### Library amplification from oligo pools

Oligo pools encoding GOIs were ordered from Twist Bioscience, and amplified by PCR using KAPA HiFi HotStart ReadyMix (Roche). To reduce biases, amplifications were monitored by real-time PCR using EvaGreen Dye (Biotium), and stopped before plateauing, as described in Klein et al., 2016 (*54*). Amplified libraries were purified by gel electrophoresis (2% agarose) and extracted using Zymoclean Gel DNA Recovery Kits (Zymo Research), followed by quantification with Qubit (Invitrogen).

### Cloning into DMX vector

Amplified libraries were cloned into the DMX vector using BsmBI-v2 NEBridge Golden Gate Assembly kit (NEB) according to the manufacturer’s protocol. The GGA product was purified using DNA Clean & Concentrator-5 (Zymo Research) quantified using Qubit (Invitrogen), and electroporated into *E. Cloni* EXPRESS BL21(DE3) electrocompetent cells (Lucigen) according to the manufacturer’s protocol. Electroporated cells were recovered at 37 °C for 1 hour in a shaking incubator at 250 rpm.

### Colony picking

Aliquots from recovered cells after electroporation were serially diluted and grown overnight at 37 °C on LB-Agar plates containing 100 μg/mL of carbenicillin; the rest of the non-diluted recovered cells were stored at 4 °C overnight. The next day, colony counting provided an estimate of CFUs in the original library, which was used to calculate the volume needed from the kept recovered cells to obtain 2,500 colonies on Bioassay plates (Corning). The cells were mixed with SOC medium and plated onto BioAssay plates containing 100 μg/ml of carbenicillin, and grown overnight at 37 °C. The next day, colonies were picked using a QPix XE Microbial Colony Picker (Molecular Devices) into ECHO-qualified 384-well plates (Beckman Coulter) containing 60 μL of low-salt LB liquid medium per well (tryptone [10 g/L], NaCl [5 g/L], yeast extract [5 g/L], carbenicillin [100 mg/L]). Plates were sealed with Breathe Easier plate seals (Fisher Scientific), and grown overnight in a shaking incubator (37 °C, 1000 rpm).

### GGA barcoding in cell lysate

Confluent cultures were transferred (2 μL) and compressed from four ECHO-qualified 384-well plates into one polypropylene 1536-well plate (Greiner) using an ECHO 525 acoustic liquid handling robot (Beckman Coulter). The 1536-well plates were sealed with Axygen® Foil Plate Seal (Fisher Scientific), inserted into a vacuum sealed bag, and heat-inactivated at 98 °C in a water bath for 30 minutes. Barcode combinations (2 ng per UMI, 8 ng in total) were transferred from a DNA barcode source plate to each well of the heat-inactivated 1536-well cell-lysate plates using the ECHO. A GGA mastermix containing BsaI-HFv2 (NEB), Salt-T4 DNA ligase (M0467L, NEB), and T4 DNA ligase buffer (NEB) was transferred to each well with the ECHO (0.5 μL). Reaction plates were incubated at 37 °C for 1 hour, followed by 5 minutes at 60 °C. The 1536-well plates were inverted and centrifuged at 200 x g to pool all the wells, and GGA products (i.e. position-specific barcodes ligated with the GOI from each well) were isolated using Zyppy Plasmid Miniprep kit (Zymo Research) according to the manufacturer’s protocol. Pools of barcoded GOIs were amplified by PCR using common primers and KAPA HiFi HotStart ReadyMix (Roche), and the product purified by gel electrophoresis (1% agarose), extracted with Zymoclean Gel DNA Recovery kit (Zymo Research), and quantified by Qubit (Invitrogen).

### Nanopore Long-read sequencing

Pools of barcoded GOIs were prepared for sequencing using Ligation Sequencing Kit V14 (SQK-LSK114, Oxford Nanopore Technologies) and loaded onto MinION Flow Cell according to the manufacturer’s protocol. Depending on library size, sequencing can be run for <72 hours.

### DMX bioinformatics pipeline

Output raw reads from ONT nanopore sequencing (.pod5) were basecalled to bam files using Dorado-0.9.1. Reads were converted and merged from BAM into Fastq files using SAMtools-1.21 (*81*) and filtered based on size and quality score using Chopper-0.9.0 (*91*) and Nanoq-0.10.0 (*92*). Filtered reads were demultiplexed into individual files based on barcode combination using Cutadapt-4.9 (*93*). Alignment to reference, and consensus sequence generation were done with Minimap2-2.28 (*94*) and samtools-1.21. A python script was used to generate a list of correct unique designs and generate pick lists to re-array ECHO culture plates into new plates for downstream processing (see Supplementary protocol DMX).

### RSV F minibinder design and screening

Design of the RSV minibinders cb4 and cb13 followed the previously published Rosetta-based design pipeline (*20*). A rotameric interaction field (RIF) was generated against the MPE8 epitope. Next, a library of 21,400 scaffolds were docked against the RIF using RifDock. An amino acid sequence was assigned to the minibinder in complex with the target using FastDesign. Designs were filtered using contact molecular surface, Rosetta ΔΔG, interface buried solvent accessible surface area and shape complementarity. Alpha-helical motifs were extracted from all designs, the motifs clustered, and the best 5,000 extracted. The scaffold library was grafted onto these motifs, designed, and filtered to improve interface quality. All designs were padded to 65 amino acids with polyserines split evenly at the N and C termini. Proteins were reverse translated using DNAWorks 2.0 (*95*). 31,394 designs were ordered as an oligo pool (Agilent).

Libraries were assembled using duplicate 25 μL reactions using Kapa HiFi polymerase (Kapa Biosystems) through qPCR (Bio-Rad, CFX96). The first reaction was used to determine half maximal amplification, which was the number of cycles used for the second amplification. The amplified DNA was then loaded onto an agarose gel and the band of the expected molecular weight excised and purified using QIAquick kits (Qiagen). This DNA was then amplified, purified using the QIA quick clean up kit (Qiagen). 6 μg of amplified DNA was transformed into yeast alongside 3 μg of linearized pETcon3 through electroporation as outlined previously (*96*).

Yeast cultures were maintained at 30 °C in C-Trp-Ura (Yeast Resource Centre, University of Washington) with 2% glucose. 18 h prior to sorting, 10 mL of SGCAA (Yeast Resource Centre, University of Washington) was inoculated with 250 μL of C-Trp-Ura culture. Cells underwent four sequential sorts. The first was to sort for expression. Cells were stained for 15 min at room temperature using 1% anti-Myc FITC (Immunology Consultants Laboratory) followed by three washes in ice cold PBS supplemented with 1 % BSA (PBSF). Immediately prior to sorting, cells were resuspended in 50 μL of PBSF. After collecting the cells that were FITC positive, these cells were grown for 24-48 h in C-Trp-Ura prior to inoculation of SGCAA. For the second round sort, cells were stained for 1 h at room temperature using 1 % anti-myc FITC, 1 μM biotinylated Ds-Cav1 (stabilized RSV F protein, (*97*)) and 250 nM of streptavidin-phycoerythrin (SAPE) (Thermo Fisher), followed by three washes in ice cold PBSF. Cells were resuspended in 50 μL of PBSF immediately prior to sorting and then the double positive (FITC and PE) population was collected. This same sort was repeated a second time. For the final sort, cells were stained with biotinylated Ds-Cav1 and SAPE in a 1:1 ratio alongside 1% anti-Myc FITC. The dilution series started at 1 μM and went for 4 steps with 3-fold dilution. The concentration of the target that achieved 50% of the saturating signal was determined using published tools (*20*).

The top hits identified from this library were then ordered again as a site saturation mutagenesis library. The sorting process above was repeated to determine affinity enhancing mutations for the designs. These most promising mutations were then recombined in a combination library using IDT ultramers and these combinatorial libraries were again sorted on yeast, as above. For the sort in the titration series that had the lowest concentration of antigen for which there was visible signal, cells were collected and streaked onto a C-Trp-Ura plate and then Sanger sequenced to identify sequence of the design. The sequences for cb4 and cb13 are:

`cb4: TLEKQLENLVRSAEVLVRHNDRLNAFFILDQAKDLAERLNDPEKLRELKELEEEL`

`cb13: GLEKQLENLVRTAEVLVRHNDRLNAFFILEQAKDLAERLNDPETLREVKELHKEL`

### RSV F cryoEM

#### CryoEM sample preparation

The stabilized prefusion RSV F ectodomain (SC-DM) was incubated with a 3-fold molar excess of the designed minibinder cb13 for 30 min at room temperature in 150 mM NaCl, 25 mM Tris pH 7.5. Complexes were applied to glow-discharged C-flat R 2/2 300-mesh copper grids and vitrified using a ThermoFisher Vitrobot Mark IV (22 °C, 100% humidity, blot time 6.5 s, blot force 0).

#### CryoEM data collection

Grids were imaged on a ThermoFisher Glacios operating at 200 kV. A total of 2,855 movies were recorded in super-resolution mode on a Falcon 4i detector, yielding a final pixel size of 0.4425 Å. Data were collected in counting mode using SerialEM with a defocus range of −1.0 to −2.0 µm. The total accumulated dose was 50 e−/Å².

#### CryoEM data processing

All processing was performed in cryoSPARC. Movies were motion-corrected and dose-weighted with Patch Motion (default parameters), and CTF parameters were estimated with Patch CTF. Initial particle picking was performed with the blob picker (diameter 160–260 Å), followed by 2D classification. The best classes were used for template picking across the dataset. Ab initio reconstructions in C1 symmetry produced well-defined maps resembling RSV-F bound to cb13. Non-uniform refinement with C1 symmetry confirmed a maximum of one cb13 minibinder per RSV-F trimer.

A local refinement with a mask excluding most of the F-protein stalk improved the global resolution from 4.60 Å to 4.53 Å. To further enhance cb13 density, 3D classification focused on the minibinder was performed into three classes (filter resolution 8 Å, O-EM batch size 5,000, learning rate 0.8, hard classification enabled). Particles enriched for bound cb13 were refined locally, yielding a final reconstruction at 4.63 Å with clearly traceable backbone density for both RSV-F and cb13. The final reconstruction data was deposited in the EMDB under accession code EMD-72769.

#### Model building and refinement

The prefusion RSV F SC-DM model was docked into the final cryoEM map and relaxed using ISOLDE. The computational design model of cb13 bound to the RSV site III epitope was similarly relaxed into the density. Due to the limited resolution, all side chains were trimmed in Phenix to Cβ atoms to avoid overinterpretation. The final cryoEM model exhibited a Cα backbone RMSD of 1.9 Å relative to the cb13 design model. The final built model was deposited in the PDB under accession code 9YCF.

### Oligomer library construction

Oligomerization domains were taken from previously validated de novo oligomerization domains (validated through EM, crystallography or SEC-MALS) (*5*, *42*, *46*). Amino acid sequences were reverse translated into DNA sequences using the tools outlined in DNA fragment generation. This DNA fragment was then ordered as a single oligo encoding ccdB at the 5’ end of the oligomer domain for vectors where the binding domain would be on the N terminus of the oligomer or 3’ where the binding domain would be on the C terminus of the oligomer domain. These were cloned into a golden gate compatible expression vector, cwby0004 (SI Table 1, Addgene ID 232213), using PaqCI with 5’ TATG and 3’ TAAT overhangs. The entry vectors encoded a GSG linker between the oligomerization domain and binding domain as well as C-terminal SNAC and 6xhis tags, mirroring the construction of LM0627. The entry vector sequences were confirmed through Sanger sequencing of the expression cassette. Cloning of the binding domains into each of the vectors then followed the same process as outlined above under “Cloning”, but in this case, the master mix contained the DNA oligo encoding the binding domain of interest, and vectors were transferred through acoustic liquid handling into each individual reaction.

### RSV neutralization

RSV neutralization assays were performed as previously described (*98*). In brief, HEp-2 cells (ATCC: CCL-23) were diluted in culture media (FluoroBrite DMEM with 10% fetal calf serum) and cells were seeded in black 384-well optically clear bottom plates (ThermoFisher Scientific) at a density of 6×10^3^ cells per well and incubated overnight at 37 °C and 5% CO_2_. The next day, serial 3-fold dilutions in culture media were performed on minibinder samples and the control MPE8 antibody in 96-well plates. Recombinant RSV A2 mKate was diluted 1:4 in culture media and equal volumes of virus dilution were added to the sample dilutions. The sample-virus mixture was incubated at 37°C for one hour and then 50 μl of the sample-virus mixture was added to the HEp-2 cells seeded the day before in 384-well plates. The plates were incubated again at 37 °C and 5% CO_2_. Fluorescence endpoints were recorded at 26 hours using excitation at 588 nm and emission at 635 nm with bottom reading on a Varioskan LUX Multimode Microplate Reader (ThermoFisher Scientific). The data was normalized using Prism version 10 (GraphPad Software Inc.) and 50% inhibitory concentrations (IC50) were calculated using a four-parameter nonlinear regression curve fit. All samples were run in duplicate and at least three independent experiments were conducted for each sample.

### Surface plasmon resonance

Stabilized RSV F (SC-DM, (*99*)) was immobilized on a CM5 chip through amine conjugation at pH 6 using a Biacore 8K (Cytiva). After washing with 1 x HBS-EP (Cytiva), we then proceeded to single cycle kinetic analysis with 120 s injection of analyte (monomeric or oligomeric cb13) followed by 60 s of dissociation. After a 6-step injection series starting at 320 pM and rising 5-fold with each subsequent injection, dissociation was measured for 1200 s. Model fitting was done using the Biacore 8K Evaluation software (Cytiva) with a Langmuir 1:1 interaction model used to fit the cb13 data and a heterogeneous ligand model used to fit the oligomer data.

## Data and code availability

Scripts and notebooks can be found at https://github.com/bwicky/SAPP_DMX. A permanent archive of the code and data will be made available at Zenodo upon final publication. The density and model of cb13 bound to RSV F is available at EMD-72769 and PDB: 9YCF. Plasmids for SAPP pipeline and oligomer screening are available through Addgene (IDs: XX-XX).

## Acknowledgements

1. L. Goldschmidt and P. Vecchiato for computational infrastructure, K. VanWormer and H. Nunez-Ortega for wetlab infrastructure, W. Chen for help with setting up NGS, This work was supported by a Human Frontier Science Program Cross Disciplinary Fellowship (LT000395/2020-C, to L.F.M.), an EMBO Non-Stipendiary Fellowship (ALTF 1047-2019, to L.F.M.), a Schmidt Science Fellowship to J.Q, an EMBO long-term Fellowship (ALTF 139-2018, to B.I.M.W.), a Caixa Foundation and Rafael del Pino Foundation Fellowships to M.E.G, a Washington Research Foundation Postdoctoral Fellowship to R.J.R, the Swedish Research Council (S.O.), the Open Philanthropy Project Improving Protein Design Fund (D.B.), the Howard Hughes Medical Institute (D.B.), the Audacious Project Project at the Institute for Protein Design (D.B.), the Defence Threats Reduction Agency and the Gates Foundation (D.B.). Biorender was used to make some figures.

## Contributions

Conceptualization: B.I.M.W., L.F.M., J.Q., R.J.R., A.M., and D.B.

Methodology: B.I.M.W., L.F.M., J.Q., R.J.R., and D.B.

Software: B.I.M.W., L.F.M., J.Q., and R.D.K.

Validation: B.I.M.W., L.F.M., J.Q., R.J.R., and X.L.

Formal analysis: B.I.M.W., L.F.M., J.Q., R.J.R., R.S., A.J.B., B.C. and X.L.

Investigation: B.I.M.W., L.F.M., J.Q., R.J.R., I.G., R.S., S.O., and M.E.

Data curation: B.I.M.W., L.F.M., J.Q., and A.J.B.

Writing – original draft: B.I.M.W., L.F.M., J.Q., and R.J.R.

Writing – review & editing: B.I.M.W., L.F.M., J.Q., R.J.R., and D.B.

Visualization: B.I.M.W., L.F.M., J.Q., and R.J.R. Supervision: K.L., D.B.

Project administration: B.I.M.W., L.F.M., J.Q., and D.B. Funding acquisition: B.I.M.W., L.F.M., J.Q., L.S., and D.B.

All authors reviewed the paper.

