## Supplementary Information for "Accelerating protein design by scaling experimental characterization"

### Supplementary Methods

#### Step-by-step protocol: SAPP

Components and reagents shown underlined refer to specific items listed with their associated catalogue numbers in Supplementary Table 6. Unless stated otherwise, chemicals were sourced from Millipore-Sigma or Fisher Scientific.

##### Gene fragment sequence design and ordering

Candidate protein sequences are reverse translated and codon optimized with a custom python script that uses the DNA chisel library to optimize the reverse translated sequence for the target organism (e.g. e_coli, h_sapiens), and removes alternative start sites where possible. DNA sequences are also checked for synthesizability with the DNA manufacturer (IDT) using their web API, to ensure that they can be ordered. Furthermore, if the GOI is too long to be ordered as a single eBlock, two compatible DNA fragments are automatically generated by searching the sequence for an optimal GGA overhang from a set of orthogonal overhangs. Finally, cloning adapters for GGA into custom entry vectors (Supplementary Table 1) are added to the sequences. The output provided by the script is a dataframe of all DNA sequences to order, expected sequences on the target plasmid, and an ordering spreadsheet that can be uploaded to the manufacturer (IDT). DNA fragments are ordered resuspended and concentration-normalized in low EDTA TE buffer (10 mM Tris, 0.1 mM EDTA, pH 8.0) in ECHO-qualified 384-well plates.

##### GGA cloning

Target plasmids must be propagated in a *ccdB* resistant strain, here we use NEB Stable cells.
Available GGA vectors are listed in Supplementary Table 1.

**Custom GGA vectors**

If required, custom GGA entry vectors can be made using template vector LM1369. This general template has a pET backbone, and features PaqCI Type IIS restriction cloning sites for GGA assembly between the start and stop codons, allowing insertion of linear DNA fragments with arbitrary N- and/or C-terminal tags and a *ccdB* cassette to facilitate creation of customized GGA entry vectors for the cloning strategy described above. A custom GGA target vector can thus be assembled in 3 days (Day 1 GGA into LM1369 and plating, Day 2 picking clones for overnight growth, Day 3 DNA prep and sequencing).

**Preparing GGA vector stocks**

GGA vectors must be propagated in *ccdB* resistant NEB Stable *E. Coli*. In brief, we transform GGA target vector plasmid into NEB Stable according to the manufacturer's protocol, plate on LB-agar plate containing the appropriate antibiotic (e.g. Kanamycin for LM0627) and pick individual clones to inoculate 50 mL of LB media (**not** rich media such as Terrific Broth).

Cultures are grown overnight in 250 mL baffled flasks. Cells are harvested by centrifugation, and DNA is extracted with a midiprep kit, with typical yields in the range of 50 to 100 µg for LM0627, which is sufficient for 2,400 – 4,800 GGA 1µL reactions (2 µg vectors is necessary for one 96-well plate).

For the GGA reaction, a molar ratio of 1:2 (vector:insert) is required.

- For GGA using the ECHO in 1 µL total reaction volume, we provide an interactive spreadsheet calculator that takes into account all parameters to achieve an optimal reaction mixture. The recipe (for NEB enzymes) uses the following ratios:

| **Amount** | **Reagent** |
| --- | --- |
| 0.1 µL | 10x T4 Ligase Buffer |
| 1.2 Units | BsaI-HFv2 restriction enzyme |
| 40 Units | T4 DNA Ligase |
| 2-4 fmol | Target GGA vector |
| 4-8 fmol | Linear DNA insert (at 2-fold excess to GGA target vector) |
| Fill to 1 µL | Nuclease-free H2O |
| **Total reaction volume 1 µL** | |

- For GGA assembled with multichannel pipettes or individual tubes by hand at 5 µL total reaction volume we provide another interactive spreadsheet calculator. The recipe (for NEB enzymes) uses the following ratios:

| **Amount** | **Reagent** |
| --- | --- |
| 0.5 µL | 10x T4 Ligase Buffer |
| 3 Units | BsaI-HFv2 restriction enzyme |
| 100 Units | T4 DNA Ligase |
| 20-30 fmol | Target GGA vector |
| 40-60 fmol | Linear DNA insert (at twofold excess to GGA target vector) |
| Fill to 5 µL | Cloning-grade H2O |
| **Total reaction volume 5 µL** | |

Reactions are assembled into 96-well PCR Plates, and spun down briefly (1,000 x g, 30 seconds) to collect reaction mixture at the bottom of each well, covered with aluminum or transparent foil, and then incubated at 37 °C for 15 minutes, followed by 5 minutes at 60 °C to reduce background.

#### Transformation and expression

**NB:** All steps are performed under the flame.
**NB:** Competent cells must stay as cold as possible and be dispensed directly after they are thawed to avoid delays by having your equipment ready before you start pipetting (mise en place).

For transformation into the *E. coli* expression strain BL21(DE3) 1 µL of the GGA reaction mixture is used.

- If using the ECHO, the 96-well PCR Plate with 1 µL of GGA reaction is directly placed on ice.
- If using the 5 µL reaction approach 1 µL of GGA reaction is transferred to the bottom of each well of a fresh 96-well PCR plate.

Ensure that the plate and reaction mixture are ice cold before proceeding (especially considering the previous 60 °C inactivation step). For optimal cooling and ease of handling we recommend to use a compatible aluminum cooling block for the 96-well PCR Plate.
**NB:** The transformation is in small volumes, and produces a few hundred CFUs when using commercial competent cells. If using competent cells with lower transformation efficiencies (e.g. lab-made), the amount of cells must be increased accordingly.

Transfer 6 µL of competent BL21(DE3) cells into each well of the 96-well PCR plate using a precooled (stored at -20 °C) dispenser tip of 200 µL total volume on an electronic repeater pipette. Typically three 200 µL aliquots of BL21(DE3) are necessary for one 96-well plate.
Pipetting cells requires careful handling as the cell suspension releases poorly from the pipette tip, we recommend to pipette onto the side of each well aligning the pipette almost horizontally and retracting the tip orthogonal to the well so that the drop releases easily.

The 96-well PCR plate is "hand centrifuged" (rapidly moving the plate inside the aluminum block downwards) to make the competent cell drops settle at the bottom of each well mixing with the GGA reaction product. Failure to complete this step will lead to failed transformations.
The competent cells with the GGA mixture are incubated covered either with a plastic lid (e.g. from a pipette tip box) or aluminum foil on ice for 30 minutes.
The cells are then heat shocked by placing the 96-well PCR plate in a dry block heater at 42 °C for 10 seconds, and then back on ice for 2 minutes.
Subsequently using an electronic repeater 100 µL of room temperature SOC media is added to each well, covered with breathable film, and the plate incubated at 37°C shaking at 1,000 rpm on an orbital plate shaker for 1 hour.

*In the meantime:*
Prepare simplified 1L of autoinduction media for protein expression under flame as:
- TB-2 (Terrific Broth II powder from MP biomedicals) autoclaved approx. 980 mL
- 2 mL of 1 M $MgSO_4$ (sterile)

- 20 mL of 50x 5052 (sterile filtered, not autoclaved)
- antibiotic, if using Kanamycin: final 50 µg/mL, so use 1 mL of 50 mg/mL kanamycin stock for 1L of media (sterile)

Media can be stored at 4-8 °C in the refrigerator for 2-3 weeks.

Under flame, dispense 1 mL per well of autoinduction media into four 96-well round bottom deep well plates for protein expression using a 1 mL multichannel pipette. The number of expression culture plates may be increased to e.g. six if more protein produced is desired.

If glycerol stocks of the cultures are desired, transfer transformations after recovery into a 96-well round bottom deep well plate containing 1 mL LB media per well with appropriate antibiotic (typically 50 µg/mL kanamycin), cover with breathable film, and incubate for 1 h at 37 °C, shaking at 1,000 rpm on an orbital plate shaker. Then dispense 50 µL into each expression plate to be incubated for 20 h as given below. Continue growing the LB media plate overnight (12-16h) at 37 °C at 1,000 rpm on an orbital plate shaker. Prepare glycerol stocks in a 96-well receiver plate as per well: 100 µL of sterile 50% (v/v) Glycerol in $H_2O$ + 100 µL cultures in LB media (swirl pipette tips in wells to mix) and cover with aluminum foil to be stored at -80 °C.

Transfer 25 µL per well of the transformations after recovery into each well of the 96-well round bottom deep well plates for expression with 1 mL of autoinduction media. Cover plates with breathable film.

Incubate for a minimum of 20 hours (do not shorten this step, your cultures may not induce) at 37˚C on an orbital plate shaker.

Harvest bacterial pellets by centrifuging at 4,000 x g for 5 minutes, discard supernatant by confidently inverting the plates and pat them dry on clean paper towels. Cover with aluminium foil and store at -80 °C or directly proceed to purification.

#### Immobilized Metal Affinity Chromatography (IMAC)

*Prepare buffers:*

Lysis buffer is mixed from B-PER chemical lysis buffer containing: 0.1 mg/mL final Lysozyme (from 100 mg/mL stock, in sterile filtered Tris 50 mM 100 mM NaCl pH 7.5, 50% (v/v) glycerol, stored aliquoted at -20˚C), Benzonase (stock diluted as 1:10000, e.g. 4 uL in 40 mL Lysis buffer), cheaper alternative: 0.01 mg/mL final DNAseI, 1mM final PMSF (**NB**: PMSF is toxic!) (from PMSF stock of 100 mM in 2-Propanol, sterile filtered). Add PMSF last as it is not stable in aqueous buffer for long and then use the buffer immediately.

Wash Buffer: 20 mM Tris, 300 mM NaCl, 25 mM Imidazole, pH 8 (can be prepared as 10x stock)

Elution Buffer: 20 mM Tris, 300 mM NaCl, 500 mM Imidazole, pH 8.

Calculate 100 µL per 1mL of culture in well to lyse, i.e. approx. 40 mL of Lysis buffer when prepping 4 full 96-well deep well plates with 1 mL of culture per well for 96 candidates tested.

Add 100 µL of lysis buffer to each well and incubate shaking at 1,000 rpm at 37 °C for 15 minutes to resuspend and lyse pellets. Consolidate everyting into one of the four 96-well round bottom plates using a multichannel pipette.

Spin down to separate the insoluble fraction, 4,000 x g for 15 minutes.

*In the meantime prepare plates for IMAC:*
Dispense 50 µL of Ni-NTA resin into a 96-well fritted plate, by dispensing 100 µL of a 50% (v/v) resuspended slurry of the resin using a large repeater pipette. Mix the slurry often by inverting its container to prevent it from settling.
Equilibrate resin on vacuum manifold by applying 3 rounds of 400 µL of wash buffer per well, make sure to apply gentle vacuum to not fully dry and crack the resin to prevent channeling.

After centrifugation, carefully collect the 400 µL of supernatant by aspiration from the bottom of the well using a 1000 µL multichannel pipette (taking care not to disturb the insoluble pellet), and directly apply that to the Ni-NTA Resin on the fritted plate, let slowly drip with no vacuum applied for approximately 5 minutes.

Perform 4 rounds of washing the resin with 400 µL of wash buffer per well. As before, be careful not to dry or crack the resin with the vacuum.
Before eluting, blot the plate spouts on Kimwipes multiple times to remove excess wash buffer.
Stack from bottom to top: a 96-well receiver plate, a 96-well sterile filtration plate, the 96-well fritted plate with the Ni-NTA resin. Add 200 µL of elution buffer to the 96-well fritted plate resin, incubate for 5 minutes.
Spin down at 1,000 x g for 1 minute to clear all elution buffer from the resin, remove the fritted plate. Spin the sterile filtration plate at 2,000 x g for 2 minutes to filter the samples. Check by eye that all have passed through the filters, sometimes longer centrifugation times may be necessary.
The final output is the 96 samples in the elution buffer, sterile filtered into 96-well receiver plates, ready for SEC.

*Purifying from from the insoluble fraction:*
For cases where the protein is mostly present in the insoluble fraction (inclusion bodies), the following protocol can be used in addition (or instead) to the protocol described above for purification from the soluble fraction. After lysis, centrifugation and removal of the soluble fraction, add Wash Buffer containing 6 M GdmCl to the insoluble pellet (add the same volume as the soluble fraction volume you just removed). Incubate at room temperature in a shaking incubator (1000 rpm) for 30 min, followed by centrifugation (4,000 x g for 15 minutes). Pipette the supernatant (making sure not to disturb the pellet), and apply to Ni-NTA resin aliquoted in a 96-well fritted plate as described above. After letting the supernatant flow through by gravity, refold the protein directly on the column by gradualing diluting the amount of GdmCl. Using vacuum at each step, and making sure to not dry the resin, apply Wash Buffer containing progressively diluted GdmCl in steps of 400 uL; 3M, 1.5 M , 0.75 M, 0.375 M, 0.18M. Finalize

refolding by washing the resin with a Wash Buffer containing no GdmCl (3 x 400 uL). Proceed to elution as described above.

*If His-tag removal is desired:*
When using vectors including a SNAC tag ([...]G|SHHW[...], cutsite indicated by "|", e.g. LM627) the His tag can be cleaved and remain on the Ni-NTA resin, so that the eluate will only contain the protein of interest without tags. The protocol is modified as follows: After the Wash steps, add 2 more Wash steps (400 µL per well) using SNAC cleavage buffer. Blot the spouts of the fritted plate using Kimwipes. Seal the bottom of the fritted plate using parafilm: Place a fresh sheet of parafilm over a 96-well deep well plate, then push the fritted plates spout first onto the flat parafilm covering said plate to align the spouts and achieve a watertight seal. SNAC buffer with 2 mM $NiCl_2$ is added to initiate the cleaving on resin. Thoroughly dry and seal top of the wells of the fritted plate using aluminum foil, and incubate shaking overnight. After at least 16 hours of incubation at RT or 37 °C for improved efficiency, collect the flowthrough, as in the protocol above. Continue to SEC.

Ni-NTA resin can be recovered from the fritted plate by adding water to the well with a squirt bottle and rapidly inverting the plate over a collection box, e.g. an empty pipette tip box, multiple times. The collected resin slurry is then collected, concentrated and stored in a large column with a 25 µm frit (Econo column, Biorad) to be stored in 20% (v/v) EtOH until regeneration (see below).

Bulk Ni-NTA resin regeneration can be done using a vacuum line with collection bottle:
**NB:** Ni ions are a health and environmental hazard, all flowthrough using Ni buffer must be collected in separate containers and disposed of according to local regulations.

- Strip and clean resin with 5 resin bed Ni-NTA Strip Buffer
- Incubate in Strip buffer for 30 min.
- Wash away strip buffer with ultrapure H2O until pH of flowthrough is near neutral (check with pH test strips)
- Do three washes of 3x resin bed volumes of 100 mM $NiSO_4$, let flow through and collect for separate disposal.
- Incubate in $NiSO_4$
- Wash out remaining $NiSO_4$ with ultrapure water, until water runs clear, collect flowthrough for separate disposal
- Store Regenerated resin in 20% (v/v) EtOH at 4 °C

#### Size-exclusion chromatography

Size exclusion chromatography is performed with 3 mL resin bed volumes columns, Cytiva Superdex 5-150 GL either S75 (3-70kDa) or S200 (10-600 kDa) depending on expected candidate protein size. To process the number of samples presented here (routinely 192 overnight, often only limited by the fraction collector capacity) an autosampler that can automatically draw and inject hundreds of samples onto the chromatography systems is required. We present methods for two instruments that can perform this: A Cytiva Akta pure with autosampler, and an Agilent 1260 bioinert with multisampler. In both setups injections are pipelined so that the imidazole elution peak elutes in the next injection before the void volume is reached. Fractionation volumes are set to 200 µL. Before each run, columns are calibrated using chromatography standards (for S75 LMW kit, Cytiva ; for S200 HMW kit, Cytiva).

*Agilent 1260 Infinity II*

The multisampler module on the Agilent 1260 and 1290 FPLC series allows the injection of the samples automatically overnight, cooled to 4 °C. The loop volume for injection should be at least 100 µL.

Samples are injected sequentially at flow rates of 0.65 mL / minute. Maintaining pump pressures that are compatible with the S200 5-150 and S75 5-150 Superdex models with an approximate pressure drop of 40 bars over the column, respecting their maximum pressure limit of 100 bars. Maximum pressure limits are determined for each column according to the manufacturer's specifications.

The fraction collector is filled with 384-well plates (Greiner Masterblock 250 µL capacity).

*Akta pure*

The Akta pure system will require an autosampler module to inject the samples automatically. To achieve low dead volume when injecting from the autosampler, a re-wiring of the Akta flow path is required. The samples are directly injected from the autosampler onto the SEC column, which then bypasses the column valve and column pressure sensors and connects directly to the UV detector. To minimize the dead volume between the UV detector and the fractionation arm, the tubing of the fractionator is switched to "blue" i.d. 0.25 mm. This "hotwiring" strategy bypasses the column pressure sensors, instead the pump system pressure is used to calculate pressure limits.

#### Analysis

The integrated data-analysis pipeline starts by generating plasmid maps from the ordered DNA fragment sequences and specified entry vectors. The open reading frame (ORF) containing the gene of interest (GOI) is automatically identified, and used to generate the expression sequence (designed sequence plus vector-specific tags), and calculate protein parameters (molecular weight, molar extinction coefficient, isoelectric point, charge etc…). Batch-exported chromatograms from either the Agilent or Akta system are analyzed using the python packages NumPy (*100*), SciPy (*77*), and Pandas (*78*, *79*), to detect peaks in the protein absorbance signal at 280 nm and convert their retention volume to approximate molecular weight compared to calibration curve assembled from standards (LMW [S75], or HMW [S200], Cytvia). In addition, by integrating the absorbance signal to determine protein concentration in each fraction, the following information can be extracted for each design: total soluble yield, polydispersity, estimated molecular weight and aggregation state of each peak. The data is exported into a standardized open format dataframe (HDF5 (*80*)). Furthermore, an algorithm for automatically picking fractions for pooling was implemented as follows; the user chooses the number of fractions to pool, and the selection logic, which can be based either on the fraction with the largest integral, or the fraction containing a peak that is closest to the expected retention volume. The list of fractions to pool is then used to generate the robot script.

#### Robotic pooling

Wells for pooling are selected using the analysis software described above. Samples are pooled and – if desired – concentration-normalized using an OT-2 pipetting robot. The software outputs an OT-2-compatible script for pooling, and a P300 single-channel pipette is used to extract relevant fractions from collection plates, and consolidate them into a 96-well plate. If the option is selected, the robot can add the appropriate amount of buffer to each consolidated well to dilute proteins to a user-defined concentration.

#### Auxiliary protocols

*Ni-NTA resin regeneration:*

Used resin is collected in a large glass column (BioRad Econo Column) for batch regeneration. Regeneration requires usage of $NiSO_4$, which is an environmental poison and skin irritant, be sure to follow proper protection measures, and dispose of all Nickel ion contaminated ions according to local regulations. Stripping Buffer (0.5 M NaOH, 100 mM EDTA).

1. Pour stripping buffer until resin is white and all green Ni is eluted
2. Cap column, incubate resin in stripping buffer as a slurry on rotator for 30 minutes
3. Pour ultrapure $H_2O$ over resin until water on resin till pH is neutral (check with pH test strips)
4. Pour green ($NiSO_4$ 100 mM in $H_2O$) solution until the entire resin is blue again and no white spots remain. Collect flow-through as nickel Waste (toxic).
5. Rinse with ultrapure $H_2O$ until eluate runs clear, collect eluate as nickel waste (toxic).
6. Adjust resin slurry to 50% resin (v/v) in 20% EtOH (v/v) for use in SAPP protocol.
7. Store regenerated resin 20% EtOH (v/v). Store at 4˚ C.

*SEC Column cleaning*

After each run SEC columns are cleaned as follows; SEC column cleaning downflow at 0.5 mL / minute if not otherwise declared (CV = Column volume). All buffers are sterile filtered (0.2 µm).

Downflow clean:

- 1 CV ultrapure H2O
- 1 CV 0.5 M NaOH at 0.2 mL / minute
- 1 CV ultrapure H2O
- 1.1 CV 20% (v/v) EtOH in $H_2O$ to store

Regularly, if the pressure drop over the column increases with time, an upflow clean is required to rinse the top column filter.

Upflow clean on Akta using the column loop in upflow, on Agilent HPLC by manually inverting column direction:

- Upflow: 1 CV ultrapure H2O 0.25 mL / minute
- Upflow: 1 CV 0.5 M NaOH at 0.13 mL / minute
- Upflow: 1 CV ultrapure H2O 0.25 mL / minute
- 1.1 CV 20% (v/v) EtOH in H2O for storage 0.25 mL / minute

### Step-by-step protocol: DMX

Components and reagents shown underlined refer to specific items listed with their associated catalogue numbers in Supplementary Table 8. Unless stated otherwise, chemicals were sourced from Millipore-Sigma or Fisher Scientific.

UMI and primers sequences can be found in Supplementary Table 3 and 4.

Schematic of library entry into the multi-purpose DMX vector is shown in Supplementary Fig. 4.

#### Day 1: Library amplification from oligo pool, and cloning into DMX vector

*1.1) Library amplification from oligo pool*

The library is amplified from an oligo pool by PCR (25 μL) with corresponding forward and reverse primers (containing BsmBI cut sites: CGTCTCgagga at 5' end and ttccgGAGACG at 3' end) (Supplementary Table 4) for 14 cycles.

| **Amount** | **Reagent** |
|---|---|
| 12.5 μL | 2x KAPA HiFi HotStart Ready Mix |
| 1.25 μL | 20x EvaGreen |
| 0.75 μL | Forward library primer (10 uM) |
| 0.75 μL | Reverse library primer (10 uM) |
| 2.5 ng | Oligo library DNA |
| Fill to 25 μL | Nuclease-free $H2O$ |

| **PCR program** | **Time** |
|---|---|
| 95°C | 3 minutes |
| 98°C | 20 sec \| 14 cycles |
| 65°C | 15 sec \| |
| 72°C | 40 sec \| |
| 72°C | 1 minute |

| | |
|---|---|
| 4°C | Hold |

The amplified library is run on a 2% agarose gel, and the corresponding band extracted with Zymoclean Gel DNA Recovery Kits (D4007/D4008, Zymo Research), eluted into 8 µL of elution buffer; 1 µL of the eluant was quantified using Qubit (Invitrogen) with dsDNA Quantitation, High Sensitivity (Q32851, Invitrogen). Expect the yield after gel extraction to be ~100 - 200 ng per PCR reaction.

*1.2) Cloning into DMX vector*

The amplified library is cloned into the DMX vector by GGA (40 uL reaction). A molar ratio of 3:1 (library:DMX vector) is used to calculate the amount of amplified library to add into the GGA reaction.

| **Amount** | **Reagent** |
|---|---|
| 40 Units | BsmBI-v2 NEBridge Golden Gate Assembly kit |
| 4 µL | 10x T4 DNA ligase buffer |
| 0.1 pmol total | DMX vector |
| 0.3 pmol total | Purified oligo library DNA |
| Fill to 40 µL | Nuclease-free H2O |

| **GGA program** | **Time** |
|---|---|
| 42°C | 60 minutes |
| 60°C | 5 minutes |
| 4°C | Hold |

The GGA product is purified using DNA Clean & Concentrator-5 (D4003, Zymo Research), eluted with 10 µL of nuclease-free H2O, then quantified using a Qubit (Invitrogen) with dsDNA Quantitation, High Sensitivity.

The GGA cloned library (90 ng) is electroporated into 50 µL of *E. Cloni* EXPRESS BL21(DE3) electrocompetent cells (Lucigen) according to the manufacturer's protocol. The cells are recovered in 1 mL in Recovery Medium at 37 °C for 1 hour in a shaking incubator at 250 rpm. **NB:** Expect the time constant from the electroporation to be ~ 4.1 ms.

Recovered cells were serially diluted (1:12,500, 1:25,000, 1:50,000) and plated on round 10-cm LB-Agar dishes containing 100 µg/mL of carbenicillin and grown overnight at 37 °C; the rest of the non-diluted recovered cells were stored at 4 °C overnight for platting the next day.

##### Day 2: CFU counting and colony generation

The number of colonies are counted from each dilution plate, and multiplied by the dilution factor to calculate the CFUs in the original library. The electroporation efficiency (CFU x library complexity) should >300 to ensure adequate coverage.

The optimal colony picking density for a 25-cm Square BioAssay agar plate (431111, Corning) is ~2,500 colonies. The volume needed from the recovered cells, previously stored 4 °C, is calculated based on the CFU, then were mixed with SOC medium to a final volume of 500 µL, which is plated using beads onto a BioAssay plate containing 100 µg/ml of carbenicillin, and grown overnight at 37 °C. **NB:** For home-made BioAssay plates, it is important to dry the agar plates sufficiently (~30 minutes) to avoid smearing of grown colonies.

##### Day 3: Colony picking

Colonies are picked using a QPix XE Microbial Colony Picker (Molecular Devices) into ECHO-qualified 384-well plates (#c74290, Beckman Coulter) containing 60 µL of low-salt LB liquid medium and 100 µg/ml of carbenicillin per well.

| **Amount** | **Low-salt LB media** |
|---|---|
| 10 g | Tryptone |
| 5 g | NaCl |
| 5 g | Yeast Extract |
| Fill to 1 L | H2O |

After picking, the plates are sealed with Breathe Easier plate seals (NC1664397, Fisher Scientific), and grown at 37 °C overnight in a shaking incubator at 1,000 rpm.

If a colony picker is not available, colonies can be manually picked into ECHO-qualified 384-well plates for demultiplexing smaller libraries (<500 genes).

##### Day 4: GGA barcoding and Nanopore sequencing

*4.1) GGA barcoding in cell-lysate*

Cells (2 µL) from confluent culture plates are transferred and compressed from four ECHO-qualified 384-well plates and into one polypropylene 1536-well plate (782270, Greiner) plate using an ECHO 525 acoustic liquid handling robot (Beckman Coulter). **NB:** prior to transfer with ECHO, it is important to invert the 384-well cultures plates for 30 minutes to allow cells to gather at the meniscus to maximize cell transfer. After the ECHO transfer, the 1536-well plates are sealed with Axygen® Foil Plate Seal (PCR-AS-600, Fisher) and inserted into a vacuum sealed bag then heat-lysed in a 98 °C water bath for 30 minutes.

A custom python script was used to: 1) randomly generate the four position-specific DNA barcode combinations and calculate their transfer volume; 2) generate the $H_2O$ transfer volume (0.5 µL - volume of barcodes) (see Jupyter notebook in DMX github repository). The ECHO is used to transfer the $H_2O$ volume followed by barcode combinations (2 ng of each of the 4 UMI for a total of 8 ng) into each well of the heat-inactivated 1536-well plates.

GGA mastermix volume (0.5 µL / well) is calculated for all the 1536-well plates and transferred using the ECHO into each well.

| **Amount** | **GGA mastermix reagents for one well** |
|---|---|
| 0.4 µL | 10x T4 Ligase Buffer |
| 1.2 Units | BsaI-HFv2 restriction enzyme |
| 40 Units | Salt-T4 DNA Ligase |
| Fill to 0.5 µL | Nuclease-free H2O |

The total volume in each well is 3.5 µL (i.e. cell lysate, barcodes, GGA mastermix).

| **Amount** | **Reagent for one well** |
|---|---|
| 2 µL | Cell-lysate |
| 0.5 µL | Barcode combination (2 ng / UMI) |
| 0.5 µL | Nuclease-free H2O |
| 0.5 µL | GGA mastermix |
| **Total reaction volume 3.5 µL** | |

| **GGA program** | **Time** |
|---|---|
| 37°C | 60 minutes |

| | |
|---|---|
| 60°C | 5 minutes |
| 4°C | Hold |

After GGA reaction, the 1536-well plates are inverted and centrifuged at 200 x g into a reservoir to pool all GGA products (i.e. 4 position-specific barcodes ligated with the GOI from each well). The pooled GGA products are purified using Zyppy Plasmid Miniprep (D4036, Zymo Research) according to the manufacturer's protocol, eluted with 20 µL of elution buffer, then quantified using a Qubit (Invitrogen). **NB:** Given that the amount of pooled DNA from the plates may saturate a Zyppy column, it is recommended to split the DNA purification across multiple columns (2 columns per 1536-well plates).

Purified DNA is amplified with the forward/reverse primer pairs (DMX1/DMX2, DMX3/DMX4, DMX5/DMX6) (Supplementary Table 4) in separate PCR reactions (25 µL).

| **Amount** | **Reagent** |
|---|---|
| 12.5 µL | 2x KAPA HiFi HotStart Ready Mix |
| 1.25 µL | Forward primer (10 uM) |
| 1.25 µL | Reverse primer (10 uM) |
| 20 ng | Purified DNA |
| Fill to 25 µL | Nuclease-free $H_2O$ |

| **PCR program** | **Time** |
|---|---|
| 95°C | 3 minutes |
| 98°C | 20 sec \| 10 cycles |
| 65°C | 15 sec \| |
| 72°C | 40 sec \| |
| 72°C | 1 minute |
| 4°C | Hold |

The amplified bands from the different PCR reactions are purified by gel electrophoresis (1% agarose) and extracted using Zymoclean Gel DNA Recovery Kits (D4007/D4008, Zymo

Research), eluted with 9 µL of elution buffer, pooled together, then quantified using a Qubit (Invitrogen) with dsDNA Quantitation, High Sensitivity (Q32851, Invitrogen).

*4.2) Nanopore long-read sequencing*

Sequencing libraries were prepared from amplified barcoded pools (200 fmol) using Ligation Sequencing Kit V14 (SQK-LSK114, Oxford Nanopore Technologies) and loaded onto MinION Flow Cell R10.4.1 (FLO-MIN114, Oxford Nanopore Technologies) according to the manufacturer's protocol. **NB:** for smaller libraries, a Flongle (FLO-MIN114/FLO-FLG114, ONT) can be used instead.

#### Generation of consensus sequences (Supplementary Fig. 6).

Scripts and tools are available on GitHub. An example dataset and a Jupyter Notebook are provided on GitHub allowing users to run the pipeline on their own.

**Software version:**

- python (3.11.3)
- dorado (0.9.1)
- samtools (1.21) (*81*)
- chopper (0.9.0) (*91*)
- nanoq (0.10.0) (*92*)
- cutadapt (4.9) (*93*)
- minimap2 (2.28) (*94*)

**User input file requirements:**

- Nanopore sequencing reads (.pod5)
- UMI and barcode combination sequences (.csv)
- Reference library sequences (.fasta)

**Steps:**

**0) Basecalling**

- Dorado: Nanopore sequencing raw reads comprise multiple .pod5 files. Super accuracy model (sup) from dorado was used to convert .pod5 files into .bam files.

**1) Read Quality filtering**

- Samtools: merge all the .bam files into one, then convert the .bam file into .fastq file.
- Chopper: combination of NanoFilt and NanoLyse that is used to filter sequences based on average read quality and length and outputs a .fastq file. We used a >Q15 cutoff and read length of 400 - 1000 bp.

- Nanoq: output summary report for nanopore reads and filtering.

**2) Barcode Demultiplexing**

- Cutadapt: demultiplex barcoded reads into individual .fastq file. The users supply lists of i) reference UMI sequences (.csv); ii) barcode combinations of 3 or 4 UMI used for each barcoded well (.csv); iii) primer pairs used for PCR amplifying the final pooled barcoded library

  # Structure of post-PCR barcoded amplicons for 4 UMI:
    - primer pair 1/2: 2-3-4-dmx7-design-dmx0-1
    - primer pair 3/4: 3-4-dmx7-design-dmx0-1-2
    - primer pair 5/6: 4-dmx7-design-dmx0-1-2-3

  # Structure of post-PCR barcoded amplicons for 3 UMI:
    - primer pair 1/2: 2-3-dmx7-design-dmx0-1
    - primer pair 3/6: 3-dmx7-design-dmx0-1-2

  Dmx0 and dmx7 are universal primers flanking design sequence.

  Cutadapt algorithm searches and trims UMI from barcoded reads that match reference UMI sequences (e.g. minimum overlap of 20 out of the full length 25 bp of each UMI), then outputs trimmed reads into file (.fastq) with the name of the well from which it originated.

**3) Alignment to Reference and Consensus Sequence Generation**

- Minimap2: aligns reads from each file to the reference gene library sequences using a modified Smith-Waterman alignment algorithm and outputs a .sam file.
- Samtools: converts .sam file into .bam file and a .bami index file, then generates consensus sequence (.fa) for each well if read depth is >150 and a base is included in the consensus if at least 51% of reads support it.

**4) Perfect Design Filtering**

- Python script: builds a list of wells with perfect sequence to re-array by filtering out any wells that 1) contain multiple consensus sequences; 2) do not perfectly match to a sequence in the reference library. Finally, if multiple wells have the same sequence, then only keep one of them.

### Supplementary Figures & Tables

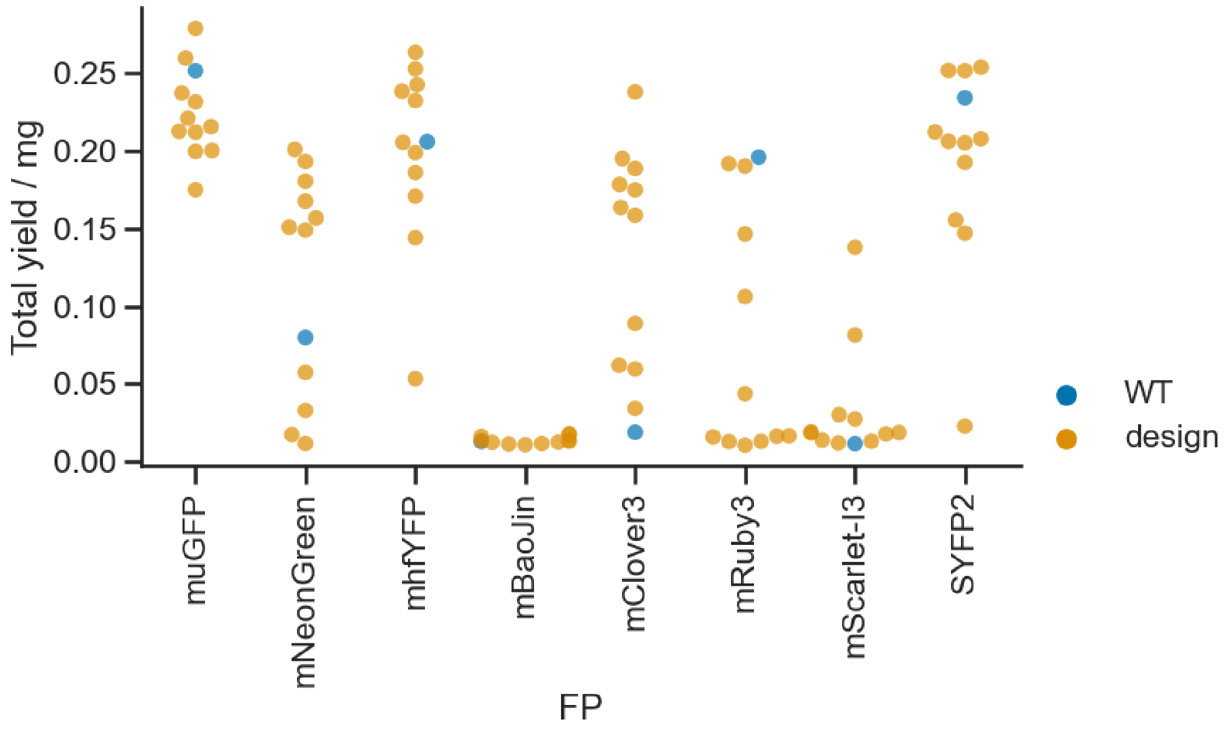

**Supplementary Fig. 1.** Total soluble yield obtained from SAPP (4 mL input culture) for FP designs, grouped by parental FP.

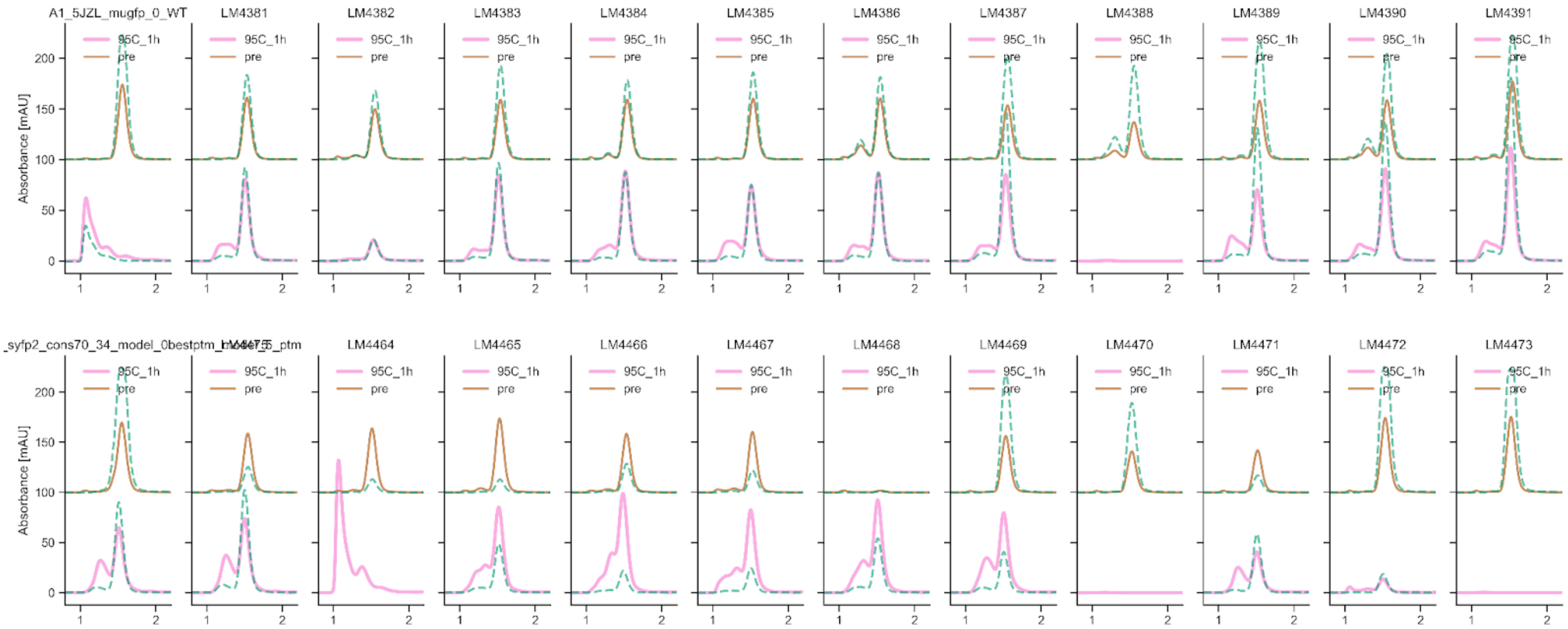

**Supplementary Fig. 2.** FP SEC chromatograms (x axes in mL) for muGFP redesigns (top) and SYFP2 redesigns (bottom) pre and post 1h incubation at 95 C. Full lines show the absorbance at 280 nm, and dashed lines show the chromophore absorbance.

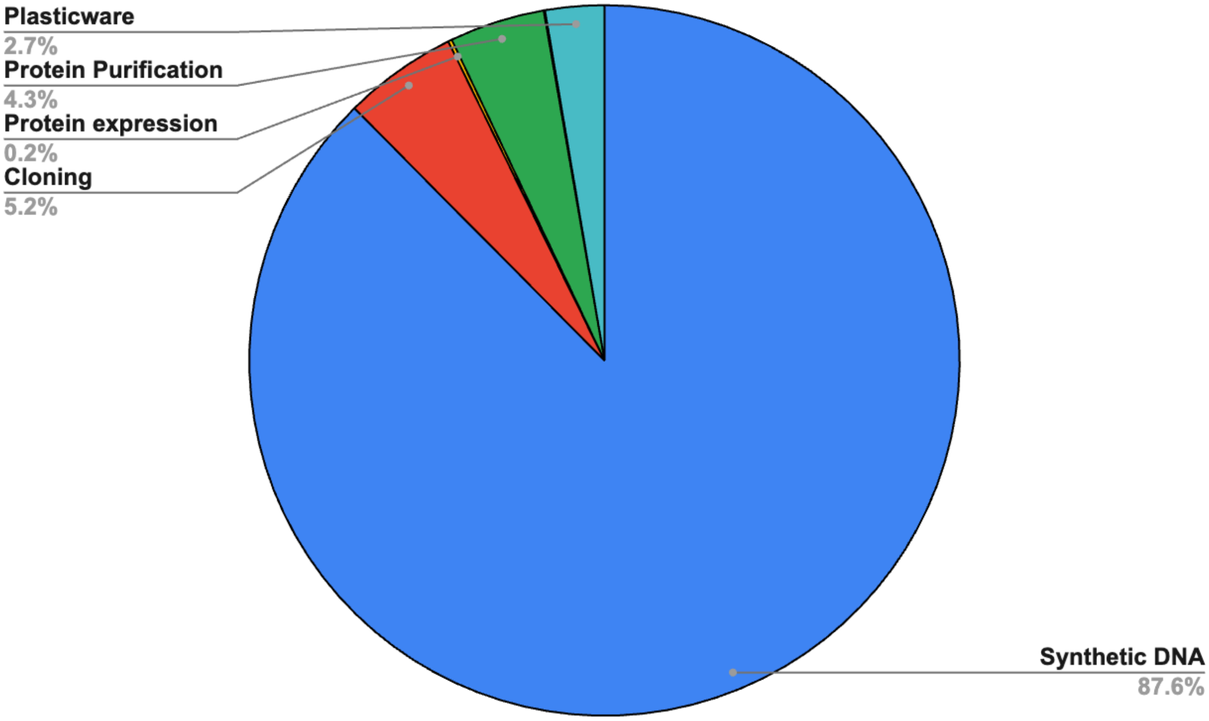

**Supplementary Fig. 3.** Cost breakdown for end-to-end execution of SAPP protocol starting from a 300 bp gene fragment (cloning/expression/purification/characterization). The total cost for one design is $24.

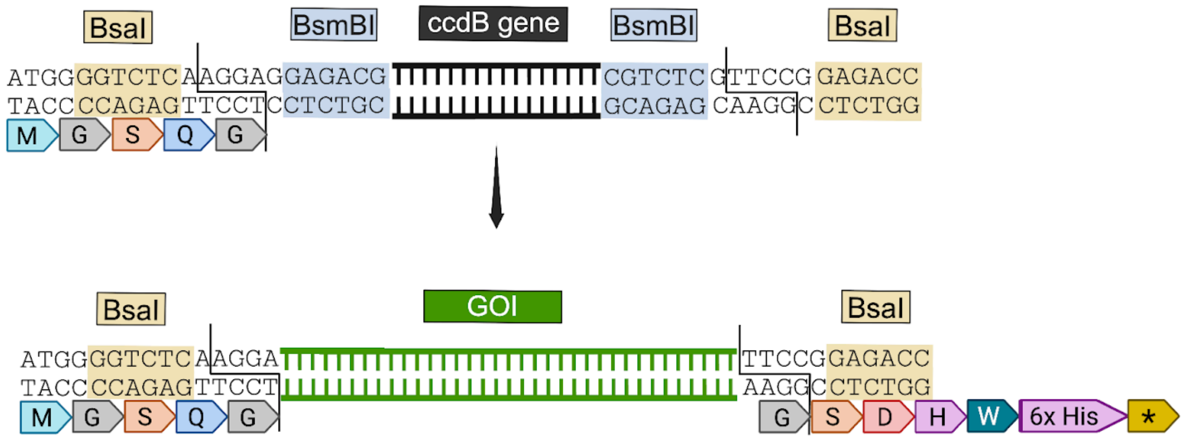

**Supplementary Fig. 4**. Multi-use cloning strategy for DMX vector. BsmBI enzyme is used to clone GOI library into the DMX vector. BsaI enzyme can be used for barcoding as well as swapping GOI with AGGA/TTCC overhangs into other SAPP entry vectors.

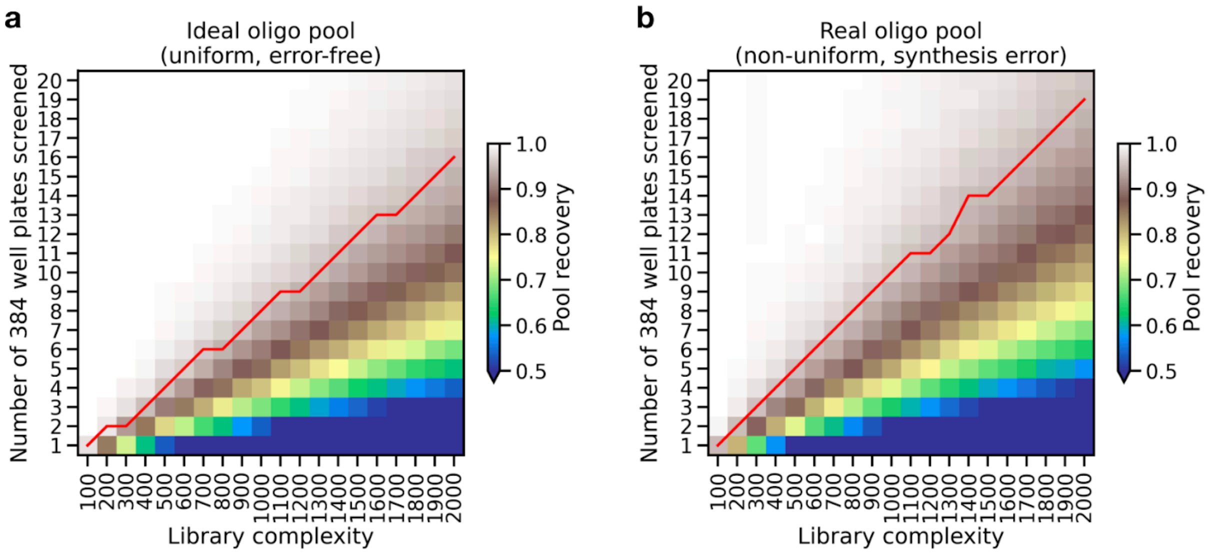

**Supplementary Fig. 5.** Simulations of expected pool recovery as a function of library complexity and number of screened colonies; (a) assuming an ideal (uniform) pool of error-free oligos, and (b) a biased pool containing oligos with synthesis error. The synthesis error rate used for the simulation was 1/3000 nt, and the oligo abundance bias assumed to be normal and approximating the distribution reported by Twist on their website. The isoline indicating 95% recovery is shown in red.

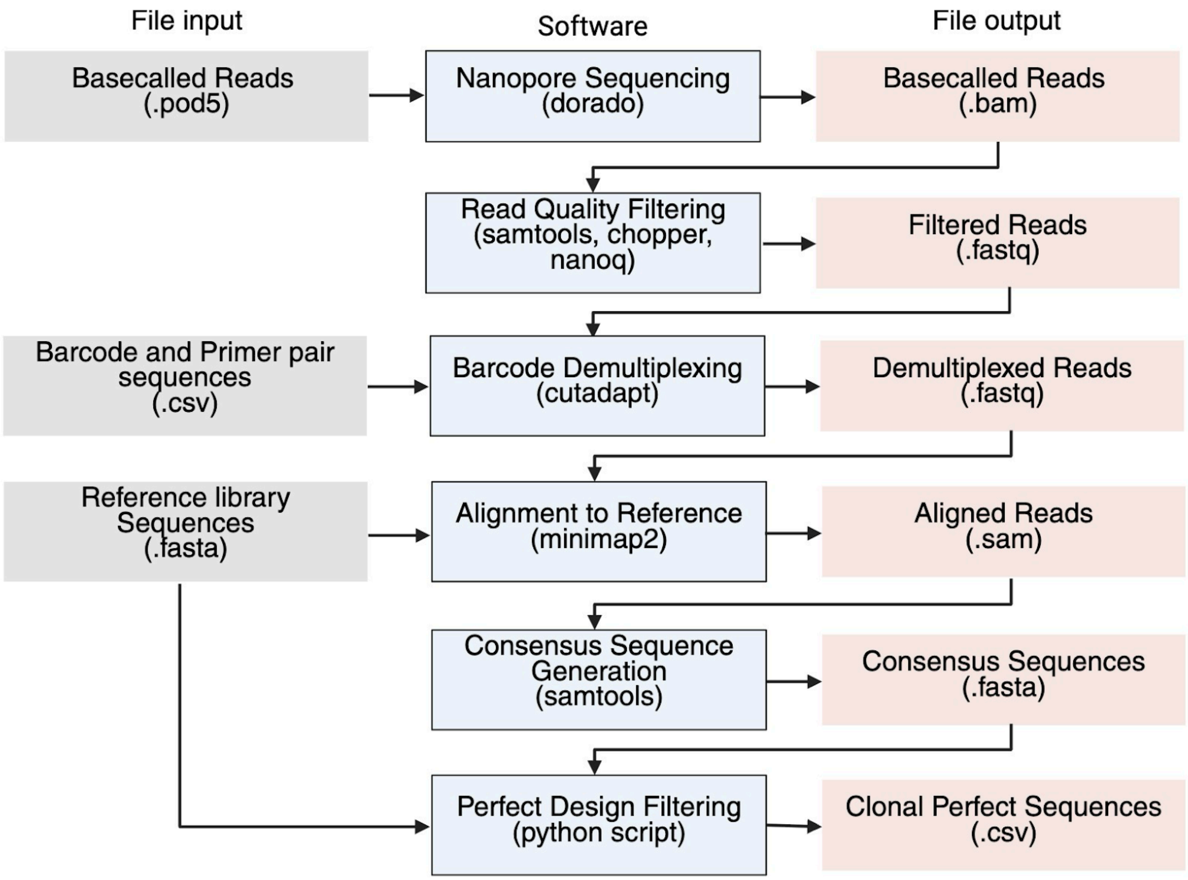

**Supplementary Fig. 6.** Workflow illustrating bioinformatic analysis to generate clonal perfect sequences from nanopore sequencing reads of pooled barcoded bacterial lysate.

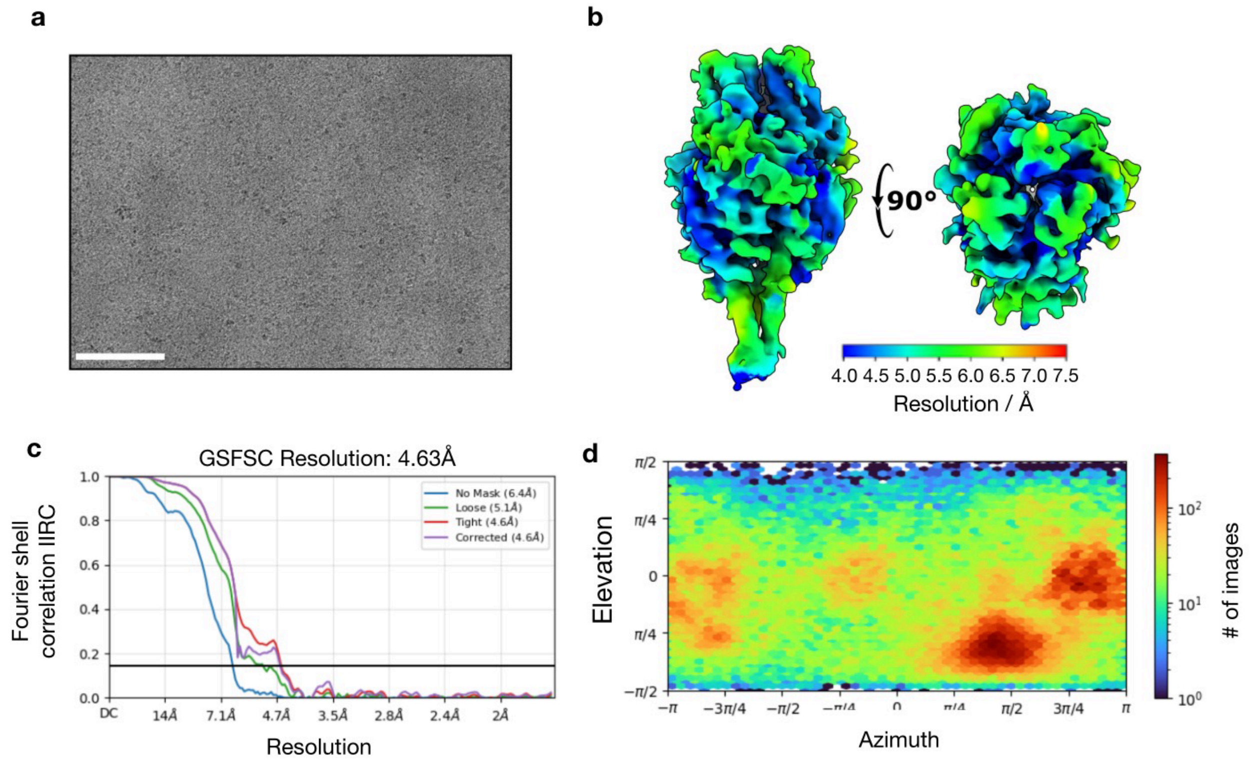

**Supplementary Fig. 7.** Cryo-EM data processing statistics for cb13 in complex with RSV F. (a) Representative raw cryo-EM micrograph of RSV F bound to cb13. Scale bar: 100 nm. (b) Local resolution map of the RSV F-cb13 complex, with values ranging from ~4 Å in the core to ~6 Å along the periphery. (c) Fourier shell correlation (FSC) curve showing the global resolution estimation with a final resolution of 4.63 Å. (d) Angular distribution plot of particle orientation used in the final 3D reconstruction.

**Supplementary Table 1.** SAPP entry vectors deposited with Addgene. All vectors are compatible with BsaI using AGGA/TTCC overhangs (exception highlighted). All plasmids have a pBR322 origin of replication (medium-copy).

| **ID** | **alternate ID** | **Description** | **Addgene ID** | **Antibiotic Resistance** | **RestrictionEnzyme** | **GGA_overhang_N** | **GGA_overhang_C** | **Link** |
|---|---|---|---|---|---|---|---|---|
| **cwby0001** | **LM627** | M-AGGA-promoter-ccdb-TTCC-SNAC-HIS | 232210 | **KanR** | BsaI | AGGA | TTCC | cwby0001 |
| **cwby0002** | **LM670** | MSG-AGGA-promoter-ccdb-TTCC-HIS | 232211 | **KanR** | BsaI | AGGA | TTCC | cwby0002 |
| **cwby0003** | **LM671** | sfGFP-AGGA-ccdb-TTCC-HIS | 232212 | **KanR** | BsaI | AGGA | TTCC | cwby0003 |
| **cwby0004** | **LM1369** | cwby template vector to insert arbitrary ccdb containing sequences: pet29-ATG-PaQCI-TAAT_GA | 232213 | **KanR** | PaqCI | TATG | TAAT | cwby0004 |
| **cwby0005** | **LM1371** | MS-his-AGGA-promoter-ccdb-TTCC | 232214 | **KanR** | BsaI | AGGA | TTCC | cwby0005 |
| **cwby0006** | **LM1423** | Cwby vector to control entire insert from N-C terminus TATG-promotor-ccdb-TAAT | 232215 | **KanR** | BsaI | TATG | TAAT | cwby0006 |
| **cwby0007** | **LM1425** | M-AGGA-promoter-ccdb-TTCC-Avitag-HIS | 232216 | **KanR** | BsaI | AGGA | TTCC | cwby0007 |
| **cwby0008** | **LM1426** | M-AGGA-promoter-ccdb-TTCC-HIS-SpyTag003 | 232217 | **KanR** | BsaI | AGGA | TTCC | cwby0008 |
| **cwby0009** | **LM1487** | his-mScarletI-AGGA-promoter-ccdb-TTCC-his | 232218 | **KanR** | BsaI | AGGA | TTCC | cwby0009 |

| | | | | | | | | |
|---|---|---|---|---|---|---|---|---|
| **cwby0010** | **LM1488** | his-TagGFP2-AGGA-promoter-ccdb-TTCC | 232219 | **KanR** | BsaI | AGGA | TTCC | cwby0010 |
| **cwby0011** | **LM1489** | his-mTagBFP2-AGGA-promoter-ccdb-TTCC | 232220 | **KanR** | BsaI | AGGA | TTCC | cwby0011 |
| **cwby0012** | **LM4372** | his-mscarletI3-AGGA-promoter-ccdb-TTCC | 232221 | **KanR** | BsaI | AGGA | TTCC | cwby0012 |
| **cwby0013** | **LM4373** | his-muGFP-AGGA-promoter-ccdb-TTCC | 232222 | **KanR** | BsaI | AGGA | TTCC | cwby0013 |
| **cwby0014** | **LM4374** | his-mBaoJin-AGGA-promoter-ccdb-TTCC | 232223 | **KanR** | BsaI | AGGA | TTCC | cwby0014 |
| **cwby0015** | **LM4376** | his-mhfYFP-AGGA-promoter-ccdb-TTCC | 232224 | **KanR** | BsaI | AGGA | TTCC | cwby0015 |
| **cwby0016** | **BW1006** | His-lgBiT-AGGA-promoter-ccdb-TTCC | 232225 | **KanR** | BsaI | AGGA | TTCC | cwby0016 |
| **cwby0017** | **BW1005** | His-GB1-smBiT-AGGA-promoter-ccdb-TTCC | 232226 | **KanR** | BsaI | AGGA | TTCC | cwby0017 |
| **cwby0018** | **BW1004** | His-smBiT-AGGA-promoter-ccdb-TTCC | 232227 | **KanR** | BsaI | AGGA | TTCC | cwby0018 |
| **cwby0019** | **BW1003** | AGGA-promoter-ccdb-TTCC-lgBiT-his | 232228 | **KanR** | BsaI | AGGA | TTCC | cwby0019 |
| **cwby0020** | **BW1002** | AGGA-promoter-ccdb-TTCC-smBiT-GB1-His | 232229 | **KanR** | BsaI | AGGA | TTCC | cwby0020 |
| **cwby0021** | **BW1001** | AGGA-promoter-ccdb-TTCC-smBiT-His | 232230 | **KanR** | BsaI | AGGA | TTCC | cwby0021 |
| **cwby0022** | **BW1008** | M-AGGA-promoter-ccdb-TTCC-SNAC-HIS CarbR | 232231 | **AmpR / CarbR** | BsaI | AGGA | TTCC | cwby0022 |
| **cwby00** | **BW102** | MSG-AGGA-promoter-ccdb-TTCC- | 23223 | **AmpR /** | BsaI | AGGA | TTCC | cwby0023 |

| | | | | | | | | |
|---|---|---|---|---|---|---|---|---|
| **23** | **2** | HIS CarbR | 2 | **CarbR** | | | | |
| **cwby0024** | **BW1023** | MS-his-AGGA-promoter-ccdb-TTCC CarbR | 232233 | **AmpR / CarbR** | BsaI | AGGA | TTCC | cwby0024 |
| **cwby0025** | **BW1024** | TATG-promotor-ccdb-TAAT CarbR | 232234 | **AmpR / CarbR** | BsaI | AGGA | TTCC | cwby0025 |
| **cwby0026** | **RR001** | AGGA-promoter-ccdb-TTCC-GS-HA LC1_004-SNAC-6xhis | TBD | **KanR** | BsaI | AGGA | TTCC | |
| **cwby0027** | **RR002** | AGGA-promoter-ccdb-TTCC-GS-HA LC1_008-SNAC-6xhis | TBD | **KanR** | BsaI | AGGA | TTCC | |
| **cwby0028** | **RR003** | AGGA-promoter-ccdb-TTCC-GS-HA LC2_059-SNAC-6xhis | TBD | **KanR** | BsaI | AGGA | TTCC | |
| **cwby0029** | **RR004** | AGGA-promoter-ccdb-TTCC-GS-HA LC2_062-SNAC-6xhis | TBD | **KanR** | BsaI | AGGA | TTCC | |
| **cwby0030** | **RR005** | AGGA-promoter-ccdb-TTCC-GS-HA LC2_063-SNAC-6xhis | TBD | **KanR** | BsaI | AGGA | TTCC | |
| **cwby0031** | **RR006** | AGGA-promoter-ccdb-TTCC-GS-HA LC2_064-SNAC-6xhis | TBD | **KanR** | BsaI | AGGA | TTCC | |
| **cwby0032** | **RR007** | AGGA-promoter-ccdb-TTCC-GS-HA LC2_065-SNAC-6xhis | TBD | **KanR** | BsaI | AGGA | TTCC | |
| **cwby0033** | **RR008** | AGGA-promoter-ccdb-TTCC-GS-HA LC2_067-SNAC-6xhis | TBD | **KanR** | BsaI | AGGA | TTCC | |
| **cwby0034** | **RR009** | AGGA-promoter-ccdb-TTCC-GS-HA LC2_068-SNAC-6xhis | TBD | **KanR** | BsaI | AGGA | TTCC | |
| **cwby0035** | **RR010** | AGGA-promoter-ccdb-TTCC-GS-HA LC3_104-SNAC-6xhis | TBD | **KanR** | BsaI | AGGA | TTCC | |
| **cwby0036** | **RR011** | AGGA-promoter-ccdb-TTCC-GS-HA LC3_109-SNAC-6xhis | TBD | **KanR** | BsaI | AGGA | TTCC | |

| | | | | | | | |
|---|---|---|---|---|---|---|---|
| **cwby0037** | **RR012** | AGGA-promoter-ccdb-TTCC-GS-HALC3_110-SNAC-6xhis | TBD | **KanR** | BsaI | AGGA | TTCC |
| **cwby0038** | **RR013** | AGGA-promoter-ccdb-TTCC-GS-HALC3_114-SNAC-6xhis | TBD | **KanR** | BsaI | AGGA | TTCC |
| **cwby0039** | **RR014** | AGGA-promoter-ccdb-TTCC-GS-HALC3_118-SNAC-6xhis | TBD | **KanR** | BsaI | AGGA | TTCC |
| **cwby0040** | **RR015** | AGGA-promoter-ccdb-TTCC-GS-HALC3_919-SNAC-6xhis | TBD | **KanR** | BsaI | AGGA | TTCC |
| **cwby0041** | **RR016** | AGGA-promoter-ccdb-TTCC-GS-SB175-SNAC-6xhis | TBD | **KanR** | BsaI | AGGA | TTCC |
| **cwby0042** | **RR017** | AGGA-promoter-ccdb-TTCC-GS-HALC4_135-SNAC-6xhis | TBD | **KanR** | BsaI | AGGA | TTCC |
| **cwby0043** | **RR018** | AGGA-promoter-ccdb-TTCC-GS-HALC4_136-SNAC-6xhis | TBD | **KanR** | BsaI | AGGA | TTCC |
| **cwby0044** | **RR019** | AGGA-promoter-ccdb-TTCC-GS-HALC4_140-SNAC-6xhis | TBD | **KanR** | BsaI | AGGA | TTCC |
| **cwby0045** | **RR020** | AGGA-promoter-ccdb-TTCC-GS-HALC5_169-SNAC-6xhis | TBD | **KanR** | BsaI | AGGA | TTCC |
| **cwby0046** | **RR021** | AGGA-promoter-ccdb-TTCC-GS-HALC5_176-SNAC-6xhis | TBD | **KanR** | BsaI | AGGA | TTCC |
| **cwby0047** | **RR022** | AGGA-HALC1_004-GS-promoter-ccdb-TTCC-SNAC-6xhis | TBD | **KanR** | BsaI | AGGA | TTCC |
| **cwby0048** | **RR023** | AGGA-HALC1_008-GS-promoter-ccdb-TTCC-SNAC-6xhis | TBD | **KanR** | BsaI | AGGA | TTCC |
| **cwby0049** | **RR024** | AGGA-HALC2_059-GS-promoter-ccdb-TTCC-SNAC-6xhis | TBD | **KanR** | BsaI | AGGA | TTCC |
| **cwby00** | **RR025** | AGGA-HALC2_062-GS-promoter-cc | TBD | **KanR** | BsaI | AGGA | TTCC |

| | | | | | | | |
|---|---|---|---|---|---|---|---|
| **50** | | db-TTCC-SNAC-6xhis | | | | | |
| **cwby0051** | **RR026** | AGGA-HALC2_063-GS-promoter-ccdb-TTCC-SNAC-6xhis | TBD | **KanR** | BsaI | AGGA | TTCC |
| **cwby0052** | **RR027** | AGGA-HALC2_064-GS-promoter-ccdb-TTCC-SNAC-6xhis | TBD | **KanR** | BsaI | AGGA | TTCC |
| **cwby0053** | **RR028** | AGGA-HALC2_065-GS-promoter-ccdb-TTCC-SNAC-6xhis | TBD | **KanR** | BsaI | AGGA | TTCC |
| **cwby0054** | **RR029** | AGGA-HALC2_067-GS-promoter-ccdb-TTCC-SNAC-6xhis | TBD | **KanR** | BsaI | AGGA | TTCC |
| **cwby0055** | **RR030** | AGGA-HALC2_068-GS-promoter-ccdb-TTCC-SNAC-6xhis | TBD | **KanR** | BsaI | AGGA | TTCC |
| **cwby0056** | **RR031** | AGGA-HALC3_104-GS-promoter-ccdb-TTCC-SNAC-6xhis | TBD | **KanR** | BsaI | AGGA | TTCC |
| **cwby0057** | **RR032** | AGGA-HALC3_109-GS-promoter-ccdb-TTCC-SNAC-6xhis | TBD | **KanR** | BsaI | AGGA | TTCC |
| **cwby0058** | **RR033** | AGGA-HALC3_110-GS-promoter-ccdb-TTCC-SNAC-6xhis | TBD | **KanR** | BsaI | AGGA | TTCC |
| **cwby0059** | **RR034** | AGGA-HALC3_114-GS-promoter-ccdb-TTCC-SNAC-6xhis | TBD | **KanR** | BsaI | AGGA | TTCC |
| **cwby0060** | **RR035** | AGGA-HALC3_118-GS-promoter-ccdb-TTCC-SNAC-6xhis | TBD | **KanR** | BsaI | AGGA | TTCC |
| **cwby0061** | **RR036** | AGGA-HALC3_919-GS-promoter-ccdb-TTCC-SNAC-6xhis | TBD | **KanR** | BsaI | AGGA | TTCC |
| **cwby0062** | **RR037** | AGGA-SB175-GS-promoter-ccdb-TTCC-SNAC-6xhis | TBD | **KanR** | BsaI | AGGA | TTCC |
| **cwby0063** | **RR038** | AGGA-HALC4_135-GS-promoter-ccdb-TTCC-SNAC-6xhis | TBD | **KanR** | BsaI | AGGA | TTCC |

| | | | | | | | |
|---|---|---|---|---|---|---|---|
| **cwby0064** | **RR039** | AGGA-HALC4_136-GS-promoter-ccdb-TTCC-SNAC-6xhis | TBD | **KanR** | BsaI | AGGA | TTCC |
| **cwby0065** | **RR040** | AGGA-HALC4_140-GS-promoter-ccdb-TTCC-SNAC-6xhis | TBD | **KanR** | BsaI | AGGA | TTCC |
| **cwby0066** | **RR041** | AGGA-HALC5_169-GS-promoter-ccdb-TTCC-SNAC-6xhis | TBD | **KanR** | BsaI | AGGA | TTCC |
| **cwby0067** | **RR042** | AGGA-HALC5_176-GS-promoter-ccdb-TTCC-SNAC-6xhis | TBD | **KanR** | BsaI | AGGA | TTCC |
| **cwby0068** | **ME001** | AGGA-promoter-ccdb-TTCC-GS-C2-02802-SNAC-6xhis | TBD | **KanR** | BsaI | AGGA | TTCC |
| **cwby0069** | **ME002** | AGGA-promoter-ccdb-TTCC-GS-C4-71-8-SNAC-6xhis | TBD | **KanR** | BsaI | AGGA | TTCC |
| **cwby0070** | **ME003** | AGGA-promoter-ccdb-TTCC-GS-C4-81-4-SNAC-6xhis | TBD | **KanR** | BsaI | AGGA | TTCC |
| **cwby0071** | **ME004** | AGGA-promoter-ccdb-TTCC-GS-C6-71-9-1-SNAC-6xhis | TBD | **KanR** | BsaI | AGGA | TTCC |
| **cwby0072** | **ME005** | AGGA-promoter-ccdb-TTCC-GS-C6-79-10-SNAC-6xhis | TBD | **KanR** | BsaI | AGGA | TTCC |
| **cwby0073** | **ME006** | AGGA-promoter-ccdb-TTCC-GS-C8-71-7-1-SNAC-6xhis | TBD | **KanR** | BsaI | AGGA | TTCC |
| **cwby0074** | **ME007** | AGGA-promoter-ccdb-TTCC-GS-H6-SNAC-6xhis | TBD | **KanR** | BsaI | AGGA | TTCC |
| **cwby0075** | **ME008** | AGGA-promoter-ccdb-TTCC-GS-H8-SNAC-6xhis | TBD | **KanR** | BsaI | AGGA | TTCC |
| **cwby0076** | **ME009** | AGGA-C2-02802-GS-promoter-ccdb-TTCC-SNAC-6xhis | TBD | **KanR** | BsaI | AGGA | TTCC |
| **cwby00** | **ME010** | AGGA-C4-71-8-GS-promoter-ccdb- | TBD | **KanR** | BsaI | AGGA | TTCC |

| | | | | | | | |
|---|---|---|---|---|---|---|---|
| **77** | | TTCC-SNAC-6xhis | | | | | |
| **cwby0078** | **ME011** | AGGA-C4-81-4-GS-promoter-ccdb-TTCC-SNAC-6xhis | TBD | **KanR** | BsaI | AGGA | TTCC |
| **cwby0079** | **ME012** | AGGA-C6-71-9-1-GS-promoter-ccdb-TTCC-SNAC-6xhis | TBD | **KanR** | BsaI | AGGA | TTCC |
| **cwby0080** | **ME013** | AGGA-C6-79-10-GS-promoter-ccdb-TTCC-SNAC-6xhis | TBD | **KanR** | BsaI | AGGA | TTCC |
| **cwby0081** | **ME014** | AGGA-C8-71-7-1-GS-promoter-ccdb-TTCC-SNAC-6xhis | TBD | **KanR** | BsaI | AGGA | TTCC |
| **cwby0082** | **ME015** | AGGA-H6-GS-promoter-ccdb-TTCC-SNAC-6xhis | TBD | **KanR** | BsaI | AGGA | TTCC |
| **cwby0083** | **ME016** | AGGA-H8-GS-promoter-ccdb-TTCC-SNAC-6xhis | TBD | **KanR** | BsaI | AGGA | TTCC |
| **DMX0001** | | M-BsaI_AGGA_BsmBI-promoter-ccdb-BsmBI_TTCC_BsaI_his-W__AMPR | TBD | **AmpR** | BsmBI/BsaI | AGGA | TTCC |
| **DMX0002** | | M-BsaI_AGGA_BsmBI-promoter-ccdb-BsmBI_TTCC_BsaI_W-his__AMPR | TBD | **AmpR** | BsmBI/BsaI | AGGA | TTCC |

**Supplementary Table 2.** Cost breakdown for SAPP protocol (starting from gene fragments).

| **Synthetic DNA** | $ / Design | Amount | Units | Usage | $ / Unit | Bulk Amount | Bulk List Price | Supplier | Cat# |
|---|---|---|---|---|---|---|---|---|---|
| eBlocks™ Gene Fragments | 21.00 | 300 | bp | 1 | $ 0.070 | 1 | $ 0.07 | IDT | N/A |
| subtotal: | 21.00 | | | | | | | | |
| **Cloning** | | | | | | | | | |
| BsaI-HF®v2 | 0.08 | 1.2 | U | 1 | $ 0.07 | 5000 | $ 331.00 | NEB | R3733L |
| T4 DNA Ligase | 0.11 | 40 | U | 1 | $ 0.00 | 100000 | $ 281.00 | NEB | M0202L |
| Entry vector (e.g. LM0627, Midiprep) | 0.13 | 14 | ng | 1 | $ 0.01 | 60000 | $ 566.00 | ZymoResearch | D4201 |
| BL21(DE3) Competent E. Coli | 0.92 | 6 | uL | 1 | $ 0.15 | 1200 | $ 184.00 | NEB | C2527I |
| subtotal: | 1.24 | | | | | | | | |
| **Protein Expression** | | | | | | | | | |
| Terrific broth II | 0.04 | 0.2 | g | 1 | $ 0.18 | 15000 | $ 2,745.10 | MP Biomedicals | 113046032 |
| Glycerol | 0.00 | 0.016 | mL | 1 | $ 0.13 | 4000 | $ 510.00 | MilliporeSigma | G5516-4L |
| Dextrose (D-Glucose), Anhydrous | 0.00 | 0.002 | g | 1 | $ 0.34 | 1000 | $ 343.50 | FisherScientific | D16-1 |
| D-Lactose monohydrate | 0.00 | 0.008 | g | 1 | $ 0.07 | 1000 | $ 72.80 | MilliporeSigma | 61345-1KG |
| Kanamycin sulfate | 0.00 | 0.0002 | g | 1 | $ 9.84 | 100 | $ 983.50 | FisherScientific | AC450811000 |
| subtotal: | 0.04 | | | | | | | | |
| **Protein Purification** | | | | | | | | | |
| B-PER™ | 0.35 | 0.4 | mL | 1 | $ 0.87 | 500 | $ 436.00 | ThermoFisher | 78248 |
| Benzonase® Nuclease | 0.12 | 10 | U | 1 | $ 0.01 | 25000 | $ 301.00 | MilliporeSigma | E1014-25KU |
| PMSF | 0.00 | 0.0001 | g | 1 | $ 12.80 | 25 | $ 320.00 | MilliporeSigma | 11359061001 |

| | | | | | | | | | |
|---|---|---|---|---|---|---|---|---|---|
| Lysozyme from chicken egg white | 0.00 | 0.00004 | g | 1 | $ 26.60 | 100 | $ 2,660.00 | MilliporeSigma | L6876 |
| HisPur™ Ni-NTA Resin | 0.02 | 0.05 | mL | 20 | $ 7.94 | 500 | $ 3,970.00 | ThermoFisher | 88223 |
| Cytiva Sx 5-150 | 0.55 | 1 | run | 5000 | $ 2,745.00 | 1 | $ 2,745.00 | Cytiva | 29148722 |
| subtotal: | 1.04 | | | | | | | | |
| **Buffers** | | | | | | | | | |
| Trizma® base | 0.00 | 0.017 | g | 1 | $ 0.13 | 10000 | $ 1,300.00 | MilliporeSigma | T1503-10KG |
| NaCl | 0.00 | 0.123 | g | 1 | $ 0.04 | 2500 | $ 95.51 | FisherScientific | BP358-212 |
| Imidazole | 0.01 | 0.007 | g | 1 | $ 1.08 | 1000 | $ 1,084.00 | MilliporeSigma | 1370981000 |
| subtotal: | 0.01 | | | | | | | | |
| **Plasticware** | | | | | | | | | |
| Tips | 0.17 | 4.000 | tip | 1 | $ 0.04 | 9600 | $ 400.00 | Opentrons | 999-00013 |
| Armadillo PCR plate | 0.08 | 0.010 | plate | 1 | $ 7.64 | 25 | $ 191.00 | FisherScientific | AB2396 |
| Deepwell culture plate | 0.01 | 0.010 | plate | 10 | $ 9.22 | 60 | $ 553.00 | FisherScientific | 278752 |
| Breathable membrane | 0.05 | 0.052 | foil | 1 | $ 0.66 | 100 | $ 65.80 | MilliporeSigma | Z763624 |
| 96w fritted plate | 0.00 | 0.010 | plate | 50 | $ 19.04 | 25 | $ 476.00 | Agilent | 200967-100 |
| 96w filter plate | 0.32 | 0.010 | plate | 1 | $ 31.96 | 25 | $ 799.00 | Agilent | 203940-100 |
| 96w receiver plate | 0.13 | 0.021 | plate | 1 | $ 6.20 | 120 | $ 743.50 | ThermoFisher | 249944 |
| 384w HPLC collection plate | 0.05 | 0.042 | plate | 10 | $ 11.73 | 60 | $ 704.00 | Greiner Bio-One | 781270 |
| subtotal: | 0.64 | | | | | | | | |
| Total cost ($) / design | 23.98 | | | | | | | | |
| Total cost ($) / 96 designs | 2,302.35 | | | | | | | | |

**Supplementary Table 3.** UMIs used for DMX barcoding.

| | BsaI5 | overhang 5 | Seq5 | UMI | Seq3 | overhang3 | BsaI3 | Final_Seq_length | BC_num | Sample_num | Final_seq |
|---|---|---|---|---|---|---|---|---|---|---|---|
| 1 | GGTCTCA | TCCT | GTGGGCCATCTTCCTGCGTATCAAA | TATGAGGACGAATCTCCCGCTTATA | GTCTAAAGTATGCGTCGCGGCATGA | GAAC | CGAGACC | 97 | 1 | 1_1 | GGTCTCATCCTGTGGGCCATCTTCCTGCGTATCAAATATGAGGACGAATCTCCCGCTTATAGTCTAAAGTATGCGTCGCGGCATGAGAACCGAGACC |
| 2 | GGTCTCA | TCCT | GTGGGCCATCTTCCTGCGTATCAAA | GGTCTTGACAAACGTGTGCTTGTAC | GTCTAAAGTATGCGTCGCGGCATGA | GAAC | CGAGACC | 97 | 1 | 1_2 | GGTCTCATCCTGTGGGCCATCTTCCTGCGTATCAAAGGTCTTGACAAACGTGTGCTTGTACGTCTAAAGTATGCGTCGCGGCATGAGAACCGAGACC |
| 3 | GGTCTCA | TCCT | GTGGGCCATCTTCCTGCGTATCAAA | GTTTATCGGGCGTGGTGCTCGCATA | GTCTAAAGTATGCGTCGCGGCATGA | GAAC | CGAGACC | 97 | 1 | 1_3 | GGTCTCATCCTGTGGGCCATCTTCCTGCGTATCAAAGTTTATCGGGCGTGGTGCTCGCATAGTCTAAAGTATGCGTCGCGGCATGAGAACCGAGACC |
| 4 | GGTCTCA | TCCT | GTGGGCCATCTTCCTGCGTATCAAA | CCGATGTTGACGGACTAATCCTGAC | GTCTAAAGTATGCGTCGCGGCATGA | GAAC | CGAGACC | 97 | 1 | 1_4 | GGTCTCATCCTGTGGGCCATCTTCCTGCGTATCAAACCGATGTTGACGGACTAATCCTGACGTCTAAAGTATGCGTCGCGGCATGAGAACCGAGACC |
| 5 | GGTCTCA | TCCT | GTGGGCCATCTTCCTGCGTATCAAA | TAGTAGTTCAGACGCCGTTAAGCGC | GTCTAAAGTATGCGTCGCGGCATGA | GAAC | CGAGACC | 97 | 1 | 1_5 | GGTCTCATCCTGTGGGCCATCTTCCTGCGTATCAAATAGTAGTTCAGACGCCGTTAAGCGCGTCTAAAGTATGCGTCGCGGCATGAGAACCGAGACC |
| 6 | GGTCTCA | TCCT | GTGGGCCATCTTCCTGCGTATCAAA | CCGTACCTAGATACACTCAATTTGT | GTCTAAAGTATGCGTCGCGGCATGA | GAAC | CGAGACC | 97 | 1 | 1_6 | GGTCTCATCCTGTGGGCCATCTTCCTGCGTATCAAACCGTACCTAGATACACTCAATTTGTGTCTAAAGTATGCGTCGCGGCATGAGAACCGAGACC |
| 7 | GGTCTCA | TCCT | GTGGGCCATCTTCCTGCGTATCAAA | CTGACGTGTGAGGCGCTAGAGCATA | GTCTAAAGTATGCGTCGCGGCATGA | GAAC | CGAGACC | 97 | 1 | 1_7 | GGTCTCATCCTGTGGGCCATCTTCCTGCGTATCAAACTGACGTGTGAGGCGCTAGAGCATAGTCTAAAGTATGCGTCGCGGCATGAGAACCGAGACC |
| 8 | GGTCTCA | TCCT | GTGGGCCATCTTCCTGCGTATCAAA | GGTATGGCACGCCTAATCTGGACAC | GTCTAAAGTATGCGTCGCGGCATGA | GAAC | CGAGACC | 97 | 1 | 1_8 | GGTCTCATCCTGTGGGCCATCTTCCTGCGTATCAAAGGTATGGCACGCCTAATCTGGACACGTCTAAAGTATGCGTCGCGGCATGAGAACCGAGACC |

| | | | | | | | | | | | |
|---|---|---|---|---|---|---|---|---|---|---|---|
| 9 | GGTCTCA | TCCT | GTGGGCCATCTTCCTGCGTATCAAA | GGATGCATGATCTAGGGCCTCGTCT | GTCTAAAGTATGCGTCGCGGCATGA | GAAC | CGAGACC | 97 | 1 | 1_9 | GGTCTCATCCTGTGGGCCATCTTCCTGCGTATCAAAGGATGCATGATCTAGGGCCTCGTCTGTCTAAAGTATGCGTCGCGGCATGAGAACCGAGACC |
| 10 | GGTCTCA | TCCT | GTGGGCCATCTTCCTGCGTATCAAA | GAGGTCTTTCATGCGTATAGTCACA | GTCTAAAGTATGCGTCGCGGCATGA | GAAC | CGAGACC | 97 | 1 | 1_10 | GGTCTCATCCTGTGGGCCATCTTCCTGCGTATCAAAGAGGTCTTTCATGCGTATAGTCACAGTCTAAAGTATGCGTCGCGGCATGAGAACCGAGACC |
| 11 | GGTCTCA | TCCT | GTGGGCCATCTTCCTGCGTATCAAA | GATTCAATATGTGTCGTCTATCCTC | GTCTAAAGTATGCGTCGCGGCATGA | GAAC | CGAGACC | 97 | 1 | 1_11 | GGTCTCATCCTGTGGGCCATCTTCCTGCGTATCAAAGATTCAATATGTGTCGTCTATCCTCGTCTAAAGTATGCGTCGCGGCATGAGAACCGAGACC |
| 12 | GGTCTCA | TCCT | GTGGGCCATCTTCCTGCGTATCAAA | GGTAACTGCGCATAGTTGGCTCTAT | GTCTAAAGTATGCGTCGCGGCATGA | GAAC | CGAGACC | 97 | 1 | 1_12 | GGTCTCATCCTGTGGGCCATCTTCCTGCGTATCAAAGGTAACTGCGCATAGTTGGCTCTATGTCTAAAGTATGCGTCGCGGCATGAGAACCGAGACC |
| 13 | GGTCTCA | TCCT | GTGGGCCATCTTCCTGCGTATCAAA | GCTCTTAAAACTGGTATCACCTGAC | GTCTAAAGTATGCGTCGCGGCATGA | GAAC | CGAGACC | 97 | 1 | 1_13 | GGTCTCATCCTGTGGGCCATCTTCCTGCGTATCAAAGCTCTTAAAACTGGTATCACCTGACGTCTAAAGTATGCGTCGCGGCATGAGAACCGAGACC |
| 14 | GGTCTCA | TCCT | GTGGGCCATCTTCCTGCGTATCAAA | GGGTGGTTAGTGATTTGCCCGTCAC | GTCTAAAGTATGCGTCGCGGCATGA | GAAC | CGAGACC | 97 | 1 | 1_14 | GGTCTCATCCTGTGGGCCATCTTCCTGCGTATCAAAGGGTGGTTAGTGATTTGCCCGTCACGTCTAAAGTATGCGTCGCGGCATGAGAACCGAGACC |
| 15 | GGTCTCA | TCCT | GTGGGCCATCTTCCTGCGTATCAAA | TAGTTGGTGGGTTTCCCTACCGTGT | GTCTAAAGTATGCGTCGCGGCATGA | GAAC | CGAGACC | 97 | 1 | 1_15 | GGTCTCATCCTGTGGGCCATCTTCCTGCGTATCAAATAGTTGGTGGGTTTCCCTACCGTGTGTCTAAAGTATGCGTCGCGGCATGAGAACCGAGACC |
| 16 | GGTCTCA | TCCT | GTGGGCCATCTTCCTGCGTATCAAA | GGTACAGTAAGTGAGAATCCTCTCT | GTCTAAAGTATGCGTCGCGGCATGA | GAAC | CGAGACC | 97 | 1 | 1_16 | GGTCTCATCCTGTGGGCCATCTTCCTGCGTATCAAAGGTACAGTAAGTGAGAATCCTCTCTGTCTAAAGTATGCGTCGCGGCATGAGAACCGAGACC |
| 17 | GGTCTCA | TCCT | GTGGGCCATCTTCCTGCGTATCAAA | GGTTCTAAGTTTAGCGTAGCCGGTT | GTCTAAAGTATGCGTCGCGGCATGA | GAAC | CGAGACC | 97 | 1 | 1_17 | GGTCTCATCCTGTGGGCCATCTTCCTGCGTATCAAAGGTTCTAAGTTTAGCGTAGCCGGTTGTCTAAAGTATGCGTCGCGGCATGAGAACCGAGACC |

| | | | | | | | | | | | |
|---|---|---|---|---|---|---|---|---|---|---|---|
| 18 | GGTCTCA | TCCT | GTGGGCCATCTTCCTGCGTATCAAA | CTTTAGGTGGGTGCGATTGCCAGTT | GTCTAAAGTATGCGTCGCGGCATGA | GAAC | CGAGACC | 97 | 1 | 1_18 | GGTCTCATCCTGTGGGCCATCTTCCTGCGTATCAAACTTTAGGTGGGTGCGATTGCCAGTTGTCTAAAGTATGCGTCGCGGCATGAGAACCGAGACC |
| 19 | GGTCTCA | TCCT | GTGGGCCATCTTCCTGCGTATCAAA | TATGTTGTGCCTTACGCCTCGATTA | GTCTAAAGTATGCGTCGCGGCATGA | GAAC | CGAGACC | 97 | 1 | 1_19 | GGTCTCATCCTGTGGGCCATCTTCCTGCGTATCAAATATGTTGTGCCTTACGCCTCGATTAGTCTAAAGTATGCGTCGCGGCATGAGAACCGAGACC |
| 20 | GGTCTCA | TCCT | GTGGGCCATCTTCCTGCGTATCAAA | TTAACCGAACTGACGGCCATCAAGG | GTCTAAAGTATGCGTCGCGGCATGA | GAAC | CGAGACC | 97 | 1 | 1_20 | GGTCTCATCCTGTGGGCCATCTTCCTGCGTATCAAATTAACCGAACTGACGGCCATCAAGGGTCTAAAGTATGCGTCGCGGCATGAGAACCGAGACC |
| 21 | GGTCTCA | TCCT | GTGGGCCATCTTCCTGCGTATCAAA | GGGTACATGCGCCTTACTCCTTGTG | GTCTAAAGTATGCGTCGCGGCATGA | GAAC | CGAGACC | 97 | 1 | 1_21 | GGTCTCATCCTGTGGGCCATCTTCCTGCGTATCAAAGGGTACATGCGCCTTACTCCTTGTGGTCTAAAGTATGCGTCGCGGCATGAGAACCGAGACC |
| 22 | GGTCTCA | TCCT | GTGGGCCATCTTCCTGCGTATCAAA | TTCTATTCTAAGCCGGCGGTCATAT | GTCTAAAGTATGCGTCGCGGCATGA | GAAC | CGAGACC | 97 | 1 | 1_22 | GGTCTCATCCTGTGGGCCATCTTCCTGCGTATCAAATTCTATTCTAAGCCGGCGGTCATATGTCTAAAGTATGCGTCGCGGCATGAGAACCGAGACC |
| 23 | GGTCTCA | TCCT | GTGGGCCATCTTCCTGCGTATCAAA | AGAACTATTTCCTGGCTGTTACGCG | GTCTAAAGTATGCGTCGCGGCATGA | GAAC | CGAGACC | 97 | 1 | 1_23 | GGTCTCATCCTGTGGGCCATCTTCCTGCGTATCAAAAGAACTATTTCCTGGCTGTTACGCGGTCTAAAGTATGCGTCGCGGCATGAGAACCGAGACC |
| 24 | GGTCTCA | TCCT | GTGGGCCATCTTCCTGCGTATCAAA | TCGGTTTCAAGGATGATCCGCGCTT | GTCTAAAGTATGCGTCGCGGCATGA | GAAC | CGAGACC | 97 | 1 | 1_24 | GGTCTCATCCTGTGGGCCATCTTCCTGCGTATCAAATCGGTTTCAAGGATGATCCGCGCTTGTCTAAAGTATGCGTCGCGGCATGAGAACCGAGACC |
| 25 | GGTCTCA | GAAC | CTGGTTGGGATCAGTCGCTTAGTGC | ACTTCTTCTCGGTCGCATGAGGCTG | GAGTGGTCCTTGGGACAGTACCCAG | AAGG | CGAGACC | 97 | 2 | 2_1 | GGTCTCAGAACCTGGTTGGGATCAGTCGCTTAGTGCACTTCTTCTCGGTCGCATGAGGCTGGAGTGGTCCTTGGGACAGTACCCAGAAGGCGAGACC |
| 26 | GGTCTCA | GAAC | CTGGTTGGGATCAGTCGCTTAGTGC | GGATACATATACGCTCGTCGGGACT | GAGTGGTCCTTGGGACAGTACCCAG | AAGG | CGAGACC | 97 | 2 | 2_2 | GGTCTCAGAACCTGGTTGGGATCAGTCGCTTAGTGCGGATACATATACGCTCGTCGGGACTGAGTGGTCCTTGGGACAGTACCCAGAAGGCGAGACC |

| | | | | | | | | | | | |
|---|---|---|---|---|---|---|---|---|---|---|---|
| 27 | GGTCTCA | GAAC | CTGGTTGGGATCAGTCGCTTAGTGC | CTCAGCCTGCCTCGCTTCTGATATT | GAGTGGTCCTTGGGACAGTACCCAG | AAGG | CGAGACC | 97 | 2 | 2_3 | GGTCTCAGAACCTGGTTGGGATCAGTCGCTTAGTGCCTCAGCCTGCCTCGCTTCTGATATTGAGTGGTCCTTGGGACAGTACCCAGAAGGCGAGACC |
| 28 | GGTCTCA | GAAC | CTGGTTGGGATCAGTCGCTTAGTGC | GGTAGGGCTACTGTTATCCTCCGTC | GAGTGGTCCTTGGGACAGTACCCAG | AAGG | CGAGACC | 97 | 2 | 2_4 | GGTCTCAGAACCTGGTTGGGATCAGTCGCTTAGTGCGGTAGGGCTACTGTTATCCTCCGTCGAGTGGTCCTTGGGACAGTACCCAGAAGGCGAGACC |
| 29 | GGTCTCA | GAAC | CTGGTTGGGATCAGTCGCTTAGTGC | CGTACGGCTGGAGAGCTGTATGTGG | GAGTGGTCCTTGGGACAGTACCCAG | AAGG | CGAGACC | 97 | 2 | 2_5 | GGTCTCAGAACCTGGTTGGGATCAGTCGCTTAGTGCCGTACGGCTGGAGAGCTGTATGTGGGAGTGGTCCTTGGGACAGTACCCAGAAGGCGAGACC |
| 30 | GGTCTCA | GAAC | CTGGTTGGGATCAGTCGCTTAGTGC | ACAGGTTGTATTACTTCGCGCCTTG | GAGTGGTCCTTGGGACAGTACCCAG | AAGG | CGAGACC | 97 | 2 | 2_6 | GGTCTCAGAACCTGGTTGGGATCAGTCGCTTAGTGCACAGGTTGTATTACTTCGCGCCTTGGAGTGGTCCTTGGGACAGTACCCAGAAGGCGAGACC |
| 31 | GGTCTCA | GAAC | CTGGTTGGGATCAGTCGCTTAGTGC | CTGGGCTCATTACAAGTGTTGCATA | GAGTGGTCCTTGGGACAGTACCCAG | AAGG | CGAGACC | 97 | 2 | 2_7 | GGTCTCAGAACCTGGTTGGGATCAGTCGCTTAGTGCCTGGGCTCATTACAAGTGTTGCATAGAGTGGTCCTTGGGACAGTACCCAGAAGGCGAGACC |
| 32 | GGTCTCA | GAAC | CTGGTTGGGATCAGTCGCTTAGTGC | CTAAGTGGCGCCGATTGTTTGTCCA | GAGTGGTCCTTGGGACAGTACCCAG | AAGG | CGAGACC | 97 | 2 | 2_8 | GGTCTCAGAACCTGGTTGGGATCAGTCGCTTAGTGCCTAAGTGGCGCCGATTGTTTGTCCAGAGTGGTCCTTGGGACAGTACCCAGAAGGCGAGACC |
| 33 | GGTCTCA | GAAC | CTGGTTGGGATCAGTCGCTTAGTGC | GTATATTTGCTCCCGGCGACGAGA | GAGTGGTCCTTGGGACAGTACCCAG | AAGG | CGAGACC | 97 | 2 | 2_9 | GGTCTCAGAACCTGGTTGGGATCAGTCGCTTAGTGCGTATATTTTGCTCCCGGCGACGAGAGAGTGGTCCTTGGGACAGTACCCAGAAGGCGAGACC |
| 34 | GGTCTCA | GAAC | CTGGTTGGGATCAGTCGCTTAGTGC | GCAATTTGCGCTTGTTCGGCATAGC | GAGTGGTCCTTGGGACAGTACCCAG | AAGG | CGAGACC | 97 | 2 | 2_10 | GGTCTCAGAACCTGGTTGGGATCAGTCGCTTAGTGCGCAATTTGCGCTTGTTCGGCATAGCGAGTGGTCCTTGGGACAGTACCCAGAAGGCGAGACC |
| 35 | GGTCTCA | GAAC | CTGGTTGGGATCAGTCGCTTAGTGC | GAGTCGAATATCCACCACCGTATGG | GAGTGGTCCTTGGGACAGTACCCAG | AAGG | CGAGACC | 97 | 2 | 2_11 | GGTCTCAGAACCTGGTTGGGATCAGTCGCTTAGTGCGAGTCGAATATCCACCACCGTATGGGAGTGGTCCTTGGGACAGTACCCAGAAGGCGAGACC |

| | | | | | | | | | | | |
|---|---|---|---|---|---|---|---|---|---|---|---|
| 36 | GGTCTCA | GAAC | CTGGTTGGGATCAGTCGCTTAGTGC | TTGTGGTTTGGGTCCTCAGAGGAGA | GAGTGGTCCTTGGGACAGTACCCAG | AAGG | CGAGACC | 97 | 2 | 2_12 | GGTCTCAGAACCTGGTTGGGATCAGTCGCTTAGTGCTTGTGGTTTGGGTCCTCAGAGGAGAGAGTGGTCCTTGGGACAGTACCCAGAAGGCGAGACC |
| 37 | GGTCTCA | GAAC | CTGGTTGGGATCAGTCGCTTAGTGC | CCGGCGCAGAAGTTTGAACGAAAAG | GAGTGGTCCTTGGGACAGTACCCAG | AAGG | CGAGACC | 97 | 2 | 2_13 | GGTCTCAGAACCTGGTTGGGATCAGTCGCTTAGTGCCCGGCGCAGAAGTTTGAACGAAAAGGAGTGGTCCTTGGGACAGTACCCAGAAGGCGAGACC |
| 38 | GGTCTCA | GAAC | CTGGTTGGGATCAGTCGCTTAGTGC | ATGCACTATTTTACGTATCCCGTGC | GAGTGGTCCTTGGGACAGTACCCAG | AAGG | CGAGACC | 97 | 2 | 2_14 | GGTCTCAGAACCTGGTTGGGATCAGTCGCTTAGTGCATGCACTATTTTACGTATCCCGTGCGAGTGGTCCTTGGGACAGTACCCAGAAGGCGAGACC |
| 39 | GGTCTCA | GAAC | CTGGTTGGGATCAGTCGCTTAGTGC | GATAGGGTGACTGCTTTCGCGTACA | GAGTGGTCCTTGGGACAGTACCCAG | AAGG | CGAGACC | 97 | 2 | 2_15 | GGTCTCAGAACCTGGTTGGGATCAGTCGCTTAGTGCGATAGGGTGACTGCTTTCGCGTACAGAGTGGTCCTTGGGACAGTACCCAGAAGGCGAGACC |
| 40 | GGTCTCA | GAAC | CTGGTTGGGATCAGTCGCTTAGTGC | TATCTGGTAGACATCTCGGCACAGA | GAGTGGTCCTTGGGACAGTACCCAG | AAGG | CGAGACC | 97 | 2 | 2_16 | GGTCTCAGAACCTGGTTGGGATCAGTCGCTTAGTGCTATCTGGTAGACATCTCGGCACAGAGAGTGGTCCTTGGGACAGTACCCAGAAGGCGAGACC |
| 41 | GGTCTCA | GAAC | CTGGTTGGGATCAGTCGCTTAGTGC | TAGTTCTGGCTATACACACTTCGGG | GAGTGGTCCTTGGGACAGTACCCAG | AAGG | CGAGACC | 97 | 2 | 2_17 | GGTCTCAGAACCTGGTTGGGATCAGTCGCTTAGTGCTAGTTCTGGCTATACACACTTCGGGGAGTGGTCCTTGGGACAGTACCCAGAAGGCGAGACC |
| 42 | GGTCTCA | GAAC | CTGGTTGGGATCAGTCGCTTAGTGC | GCATAGAGTTACCCGATGGATTCGA | GAGTGGTCCTTGGGACAGTACCCAG | AAGG | CGAGACC | 97 | 2 | 2_18 | GGTCTCAGAACCTGGTTGGGATCAGTCGCTTAGTGCGCATAGAGTTACCCGATGGATTCGAGAGTGGTCCTTGGGACAGTACCCAGAAGGCGAGACC |
| 43 | GGTCTCA | GAAC | CTGGTTGGGATCAGTCGCTTAGTGC | GTTCATGGTACAGGCTTCTTTACGG | GAGTGGTCCTTGGGACAGTACCCAG | AAGG | CGAGACC | 97 | 2 | 2_19 | GGTCTCAGAACCTGGTTGGGATCAGTCGCTTAGTGCGTTCATGGTACAGGCTTCTTTACGGGAGTGGTCCTTGGGACAGTACCCAGAAGGCGAGACC |
| 44 | GGTCTCA | GAAC | CTGGTTGGGATCAGTCGCTTAGTGC | CGATCTCGGGCCTGGGTTTTGAGTA | GAGTGGTCCTTGGGACAGTACCCAG | AAGG | CGAGACC | 97 | 2 | 2_20 | GGTCTCAGAACCTGGTTGGGATCAGTCGCTTAGTGCCGATCTCGGGCCTGGGTTTTGAGTAGAGTGGTCCTTGGGACAGTACCCAGAAGGCGAGACC |

| | | | | | | | | | | | |
|---|---|---|---|---|---|---|---|---|---|---|---|
| 45 | GGTCTCA | GAAC | CTGGTTGGGATCAGTCGCTTAGTGC | ATTATTCGTGACCCAACTCATCAGG | GAGTGGTCCTTGGGACAGTACCCAG | AAGG | CGAGACC | 97 | 2 | 2_21 | GGTCTCAGAACCTGGTTGGGATCAGTCGCTTAGTGCATTATTCGTGACCCAACTCATCAGGGAGTGGTCCTTGGGACAGTACCCAGAAGGCGAGACC |
| 46 | GGTCTCA | GAAC | CTGGTTGGGATCAGTCGCTTAGTGC | CTGAATGGTGAATAATGCGTTCGCC | GAGTGGTCCTTGGGACAGTACCCAG | AAGG | CGAGACC | 97 | 2 | 2_22 | GGTCTCAGAACCTGGTTGGGATCAGTCGCTTAGTGCCTGAATGGTGAATAATGCGTTCGCCGAGTGGTCCTTGGGACAGTACCCAGAAGGCGAGACC |
| 47 | GGTCTCA | GAAC | CTGGTTGGGATCAGTCGCTTAGTGC | AGAAAGTCTTGGATACACGGCCGGG | GAGTGGTCCTTGGGACAGTACCCAG | AAGG | CGAGACC | 97 | 2 | 2_23 | GGTCTCAGAACCTGGTTGGGATCAGTCGCTTAGTGCAGAAAGTCTTGGATACACGGCCGGGGAGTGGTCCTTGGGACAGTACCCAGAAGGCGAGACC |
| 48 | GGTCTCA | GAAC | CTGGTTGGGATCAGTCGCTTAGTGC | GTGTGTTCCTATGCACAATTTCATA | GAGTGGTCCTTGGGACAGTACCCAG | AAGG | CGAGACC | 97 | 2 | 2_24 | GGTCTCAGAACCTGGTTGGGATCAGTCGCTTAGTGCGTGTGTTCCTATGCACAATTTCATAGAGTGGTCCTTGGGACAGTACCCAGAAGGCGAGACC |
| 49 | GGTCTCA | AAGG | CGACACCGAACGTGCGACAAAACTA | TCGTTTGGAGCCGTTCACACATGAA | TGGATAAACCGGCTAGGTCGCAGAG | CTGA | CGAGACC | 97 | 3 | 3_1 | GGTCTCAAAGGCGACACCGAACGTGCGACAAAACTATCGTTTGGAGCCGTTCACACATGAATGGATAAACCGGCTAGGTCGCAGAGCTGACGAGACC |
| 50 | GGTCTCA | AAGG | CGACACCGAACGTGCGACAAAACTA | CTGATCAACTTGCGCCCAGCGTTAT | TGGATAAACCGGCTAGGTCGCAGAG | CTGA | CGAGACC | 97 | 3 | 3_2 | GGTCTCAAAGGCGACACCGAACGTGCGACAAAACTACTGATCAACTTGCGCCCAGCGTTATTGGATAAACCGGCTAGGTCGCAGAGCTGACGAGACC |
| 51 | GGTCTCA | AAGG | CGACACCGAACGTGCGACAAAACTA | GACGATGTTGCCTGTTTTGATACGA | TGGATAAACCGGCTAGGTCGCAGAG | CTGA | CGAGACC | 97 | 3 | 3_3 | GGTCTCAAAGGCGACACCGAACGTGCGACAAAACTAGACGATGTTGCCTGTTTTGATACGATGGATAAACCGGCTAGGTCGCAGAGCTGACGAGACC |
| 52 | GGTCTCA | AAGG | CGACACCGAACGTGCGACAAAACTA | GGGTAGTCGTGAGGTGAACTCTTCC | TGGATAAACCGGCTAGGTCGCAGAG | CTGA | CGAGACC | 97 | 3 | 3_4 | GGTCTCAAAGGCGACACCGAACGTGCGACAAAACTAGGGTAGTCGTGAGGTGAACTCTTCCTGGATAAACCGGCTAGGTCGCAGAGCTGACGAGACC |
| 53 | GGTCTCA | AAGG | CGACACCGAACGTGCGACAAAACTA | AGCCATTTTACGATTCTATTCGATG | TGGATAAACCGGCTAGGTCGCAGAG | CTGA | CGAGACC | 97 | 3 | 3_5 | GGTCTCAAAGGCGACACCGAACGTGCGACAAAACTAAGCCATTTTACGATTCTATTCGATGTGGATAAACCGGCTAGGTCGCAGAGCTGACGAGACC |

| | | | | | | | | | | | |
|---|---|---|---|---|---|---|---|---|---|---|---|
| 54 | GGTCTCA | AAGG | CGACACCGAACGTGCGACAAAACTA | GTGGTTTATATAATCCCACCTCCTA | TGGATAAACCGGCTAGGTCGCAGAG | CTGA | CGAGACC | 97 | 3 | 3_6 | GGTCTCAAAGGCGACACCGAACGTGCGACAAAACTAGTGGTTTATATAATCCCACCTCCTATGGATAAACCGGCTAGGTCGCAGAGCTGACGAGACC |
| 55 | GGTCTCA | AAGG | CGACACCGAACGTGCGACAAAACTA | GCGAAGAACATCCCGGCATTTCATG | TGGATAAACCGGCTAGGTCGCAGAG | CTGA | CGAGACC | 97 | 3 | 3_7 | GGTCTCAAAGGCGACACCGAACGTGCGACAAAACTAGCGAAGAACATCCCGGCATTTCATGTGGATAAACCGGCTAGGTCGCAGAGCTGACGAGACC |
| 56 | GGTCTCA | AAGG | CGACACCGAACGTGCGACAAAACTA | GCTGGGACAATGCCGAAAACTCTTC | TGGATAAACCGGCTAGGTCGCAGAG | CTGA | CGAGACC | 97 | 3 | 3_8 | GGTCTCAAAGGCGACACCGAACGTGCGACAAAACTAGCTGGGACAATGCCGAAAACTCTTCTGGATAAACCGGCTAGGTCGCAGAGCTGACGAGACC |
| 57 | GGTCTCA | AAGG | CGACACCGAACGTGCGACAAAACTA | ATTCCGTACCAACCCGCGTCTTAGA | TGGATAAACCGGCTAGGTCGCAGAG | CTGA | CGAGACC | 97 | 3 | 3_9 | GGTCTCAAAGGCGACACCGAACGTGCGACAAAACTAATTCCGTACCAACCCGCGTCTTAGATGGATAAACCGGCTAGGTCGCAGAGCTGACGAGACC |
| 58 | GGTCTCA | AAGG | CGACACCGAACGTGCGACAAAACTA | GCTGAGGAAGCCCAATGTTCAGTAC | TGGATAAACCGGCTAGGTCGCAGAG | CTGA | CGAGACC | 97 | 3 | 3_10 | GGTCTCAAAGGCGACACCGAACGTGCGACAAAACTAGCTGAGGAAGCCCAATGTTCAGTACTGGATAAACCGGCTAGGTCGCAGAGCTGACGAGACC |
| 59 | GGTCTCA | AAGG | CGACACCGAACGTGCGACAAAACTA | GTAACCTTAGCACGCCGGAGTGGAG | TGGATAAACCGGCTAGGTCGCAGAG | CTGA | CGAGACC | 97 | 3 | 3_11 | GGTCTCAAAGGCGACACCGAACGTGCGACAAAACTAGTAACCTTAGCACGCCGGAGTGGAGTGGATAAACCGGCTAGGTCGCAGAGCTGACGAGACC |
| 60 | GGTCTCA | AAGG | CGACACCGAACGTGCGACAAAACTA | GCGATTAGTTCTGTTGCTAAACCAG | TGGATAAACCGGCTAGGTCGCAGAG | CTGA | CGAGACC | 97 | 3 | 3_12 | GGTCTCAAAGGCGACACCGAACGTGCGACAAAACTAGCGATTAGTTCTGTTGCTAAACCAGTGGATAAACCGGCTAGGTCGCAGAGCTGACGAGACC |
| 61 | GGTCTCA | AAGG | CGACACCGAACGTGCGACAAAACTA | CTTAAAGGTGATTCACACGTGTGCC | TGGATAAACCGGCTAGGTCGCAGAG | CTGA | CGAGACC | 97 | 3 | 3_13 | GGTCTCAAAGGCGACACCGAACGTGCGACAAAACTACTTAAAGGTGATTCACACGTGTGCCTGGATAAACCGGCTAGGTCGCAGAGCTGACGAGACC |
| 62 | GGTCTCA | AAGG | CGACACCGAACGTGCGACAAAACTA | ATACTTATTCCGCTCATTGCACAGG | TGGATAAACCGGCTAGGTCGCAGAG | CTGA | CGAGACC | 97 | 3 | 3_14 | GGTCTCAAAGGCGACACCGAACGTGCGACAAAACTAATACTTATTCCGCTCATTGCACAGGTGGATAAACCGGCTAGGTCGCAGAGCTGACGAGACC |

| | | | | | | | | | | | |
|---|---|---|---|---|---|---|---|---|---|---|---|
| 63 | GGTCTCA | AAGG | CGACACCGAACGTGCGACAAAACTA | TGAAACCAATTTCACCTCAGCGGCG | TGGATAAACCGGCTAGGTCGCAGAG | CTGA | CGAGACC | 97 | 3 | 3_15 | GGTCTCAAAGGCGACACCGAACGTGCGACAAAACTATGAAACCAATTTCACCTCAGCGGCGTGGATAAACCGGCTAGGTCGCAGAGCTGACGAGACC |
| 64 | GGTCTCA | AAGG | CGACACCGAACGTGCGACAAAACTA | CGACCACTCGCCTCCCGTTATGATC | TGGATAAACCGGCTAGGTCGCAGAG | CTGA | CGAGACC | 97 | 3 | 3_16 | GGTCTCAAAGGCGACACCGAACGTGCGACAAAACTACGACCACTCGCCTCCCGTTATGATCTGGATAAACCGGCTAGGTCGCAGAGCTGACGAGACC |
| 65 | GGTCTCA | AAGG | CGACACCGAACGTGCGACAAAACTA | GTTAATGCTGTTTAGCGTAACCTCG | TGGATAAACCGGCTAGGTCGCAGAG | CTGA | CGAGACC | 97 | 3 | 3_17 | GGTCTCAAAGGCGACACCGAACGTGCGACAAAACTAGTTAATGCTGTTTAGCGTAACCTCGTGGATAAACCGGCTAGGTCGCAGAGCTGACGAGACC |
| 66 | GGTCTCA | AAGG | CGACACCGAACGTGCGACAAAACTA | GTCTAATATGGCGTAGCTCAACGCG | TGGATAAACCGGCTAGGTCGCAGAG | CTGA | CGAGACC | 97 | 3 | 3_18 | GGTCTCAAAGGCGACACCGAACGTGCGACAAAACTAGTCTAATATGGCGTAGCTCAACGCGTGGATAAACCGGCTAGGTCGCAGAGCTGACGAGACC |
| 67 | GGTCTCA | AAGG | CGACACCGAACGTGCGACAAAACTA | AGTAAATGGCCTGACCGGAGTTGGT | TGGATAAACCGGCTAGGTCGCAGAG | CTGA | CGAGACC | 97 | 3 | 3_19 | GGTCTCAAAGGCGACACCGAACGTGCGACAAAACTAAGTAAATGGCCTGACCGGAGTTGGTTGGATAAACCGGCTAGGTCGCAGAGCTGACGAGACC |
| 68 | GGTCTCA | AAGG | CGACACCGAACGTGCGACAAAACTA | ATGTGAAATGAGGTCTACCCTGTCA | TGGATAAACCGGCTAGGTCGCAGAG | CTGA | CGAGACC | 97 | 3 | 3_20 | GGTCTCAAAGGCGACACCGAACGTGCGACAAAACTAATGTGAAATGAGGTCTACCCTGTCATGGATAAACCGGCTAGGTCGCAGAGCTGACGAGACC |
| 69 | GGTCTCA | AAGG | CGACACCGAACGTGCGACAAAACTA | GTGTGTCCTACCTACGCGAGGAATT | TGGATAAACCGGCTAGGTCGCAGAG | CTGA | CGAGACC | 97 | 3 | 3_21 | GGTCTCAAAGGCGACACCGAACGTGCGACAAAACTAGTGTGTCCTACCTACGCGAGGAATTTGGATAAACCGGCTAGGTCGCAGAGCTGACGAGACC |
| 70 | GGTCTCA | AAGG | CGACACCGAACGTGCGACAAAACTA | TTTGGACGATATACAGGACCCGGTG | TGGATAAACCGGCTAGGTCGCAGAG | CTGA | CGAGACC | 97 | 3 | 3_22 | GGTCTCAAAGGCGACACCGAACGTGCGACAAAACTATTTGGACGATATACAGGACCCGGTGTGGATAAACCGGCTAGGTCGCAGAGCTGACGAGACC |
| 71 | GGTCTCA | AAGG | CGACACCGAACGTGCGACAAAACTA | AAGCGTACCAACTACCTCCGAGTCT | TGGATAAACCGGCTAGGTCGCAGAG | CTGA | CGAGACC | 97 | 3 | 3_23 | GGTCTCAAAGGCGACACCGAACGTGCGACAAAACTAAAGCGTACCAACTACCTCCGAGTCTTGGATAAACCGGCTAGGTCGCAGAGCTGACGAGACC |

| | | | | | | | | | | | |
|---|---|---|---|---|---|---|---|---|---|---|---|
| 72 | GGTCTCA | AAGG | CGACACCGAACGTGCGACAAAACTA | TTGTGTCGGCCACAGGTACGAAAAC | TGGATAAACCGGCTAGGTCGCAGAG | CTGA | CGAGACC | 97 | 3 | 3_24 | GGTCTCAAAGGCGACACCGAACGTGCGACAAAACTATTGTGTCGGCCACAGGTACGAAAACTGGATAAACCGGCTAGGTCGCAGAGCTGACGAGACC |
| 73 | GGTCTCA | CTGA | TTTATTCAGGGCACTACCCGGAGCT | TAGCGATTATGTTCCCGTTTTAAGC | ATCGGTGACGGCGATTCTCACATTT | GGAA | CGAGACC | 97 | 4 | 4_1 | GGTCTCACTGATTTATTCAGGGCACTACCCGGAGCTTAGCGATTATGTTCCCGTTTTAAGCATCGGTGACGGCGATTCTCACATTTGGAACGAGACC |
| 74 | GGTCTCA | CTGA | TTTATTCAGGGCACTACCCGGAGCT | GCTATGGCTCGTGAAGTGAAACGAA | ATCGGTGACGGCGATTCTCACATTT | GGAA | CGAGACC | 97 | 4 | 4_2 | GGTCTCACTGATTTATTCAGGGCACTACCCGGAGCTGCTATGGCTCGTGAAGTGAAACGAAATCGGTGACGGCGATTCTCACATTTGGAACGAGACC |
| 75 | GGTCTCA | CTGA | TTTATTCAGGGCACTACCCGGAGCT | ACCGTATGGGCGTCATAAAGGGAGC | ATCGGTGACGGCGATTCTCACATTT | GGAA | CGAGACC | 97 | 4 | 4_3 | GGTCTCACTGATTTATTCAGGGCACTACCCGGAGCTACCGTATGGGCGTCATAAAGGGAGCATCGGTGACGGCGATTCTCACATTTGGAACGAGACC |
| 76 | GGTCTCA | CTGA | TTTATTCAGGGCACTACCCGGAGCT | GTAGTTTTCTCATTTCGCGCCTCGG | ATCGGTGACGGCGATTCTCACATTT | GGAA | CGAGACC | 97 | 4 | 4_4 | GGTCTCACTGATTTATTCAGGGCACTACCCGGAGCTGTAGTTTTCTCATTTCGCGCCTCGGATCGGTGACGGCGATTCTCACATTTGGAACGAGACC |
| 77 | GGTCTCA | CTGA | TTTATTCAGGGCACTACCCGGAGCT | TGGGATGGCGTCGTCTTAGCGTTAT | ATCGGTGACGGCGATTCTCACATTT | GGAA | CGAGACC | 97 | 4 | 4_5 | GGTCTCACTGATTTATTCAGGGCACTACCCGGAGCTTGGGATGGCGTCGTCTTAGCGTTATATCGGTGACGGCGATTCTCACATTTGGAACGAGACC |
| 78 | GGTCTCA | CTGA | TTTATTCAGGGCACTACCCGGAGCT | GGCGATATAGGAGAGTCCGCGCTAA | ATCGGTGACGGCGATTCTCACATTT | GGAA | CGAGACC | 97 | 4 | 4_6 | GGTCTCACTGATTTATTCAGGGCACTACCCGGAGCTGGCGATATAGGAGAGTCCGCGCTAAATCGGTGACGGCGATTCTCACATTTGGAACGAGACC |
| 79 | GGTCTCA | CTGA | TTTATTCAGGGCACTACCCGGAGCT | TTCATATTTAGGTCGTCCGCTCTTA | ATCGGTGACGGCGATTCTCACATTT | GGAA | CGAGACC | 97 | 4 | 4_7 | GGTCTCACTGATTTATTCAGGGCACTACCCGGAGCTTTCATATTTAGGTCGTCCGCTCTTAATCGGTGACGGCGATTCTCACATTTGGAACGAGACC |
| 80 | GGTCTCA | CTGA | TTTATTCAGGGCACTACCCGGAGCT | GGTCGGTCAATTCCCTCCTAAGGAG | ATCGGTGACGGCGATTCTCACATTT | GGAA | CGAGACC | 97 | 4 | 4_8 | GGTCTCACTGATTTATTCAGGGCACTACCCGGAGCTGGTCGGTCAATTCCCTCCTAAGGAGATCGGTGACGGCGATTCTCACATTTGGAACGAGACC |

| | | | | | | | | | | | |
|---|---|---|---|---|---|---|---|---|---|---|---|
| 81 | GGTCTCA | CTGA | TTTATTCAGGGCACTACCCGGAGCT | GCTAACTGGAACGTCCCGGAATTGC | ATCGGTGACGGCGATTCTCACATTT | GGAA | CGAGACC | 97 | 4 | 4_9 | GGTCTCACTGATTTATTCAGGGCACTACCCGGAGCTGCTAACTGGAACGTCCCGGAATTGCATCGGTGACGGCGATTCTCACATTTGGAACGAGACC |
| 82 | GGTCTCA | CTGA | TTTATTCAGGGCACTACCCGGAGCT | AGTCTTTCACAACCCTACGGTGACT | ATCGGTGACGGCGATTCTCACATTT | GGAA | CGAGACC | 97 | 4 | 4_10 | GGTCTCACTGATTTATTCAGGGCACTACCCGGAGCTAGTCTTTCACAACCCTACGGTGACTATCGGTGACGGCGATTCTCACATTTGGAACGAGACC |
| 83 | GGTCTCA | CTGA | TTTATTCAGGGCACTACCCGGAGCT | GGTGACTTATGAACCTTTGCGCATT | ATCGGTGACGGCGATTCTCACATTT | GGAA | CGAGACC | 97 | 4 | 4_11 | GGTCTCACTGATTTATTCAGGGCACTACCCGGAGCTGGTGACTTATGAACCTTTGCGCATTATCGGTGACGGCGATTCTCACATTTGGAACGAGACC |
| 84 | GGTCTCA | CTGA | TTTATTCAGGGCACTACCCGGAGCT | TATTCGAGGATGGATTCGGGCTCCT | ATCGGTGACGGCGATTCTCACATTT | GGAA | CGAGACC | 97 | 4 | 4_12 | GGTCTCACTGATTTATTCAGGGCACTACCCGGAGCTTATTCGAGGATGGATTCGGGCTCCTATCGGTGACGGCGATTCTCACATTTGGAACGAGACC |
| 85 | GGTCTCA | CTGA | TTTATTCAGGGCACTACCCGGAGCT | TCATTCGCTAAATCAGTCGAGTTAG | ATCGGTGACGGCGATTCTCACATTT | GGAA | CGAGACC | 97 | 4 | 4_13 | GGTCTCACTGATTTATTCAGGGCACTACCCGGAGCTTCATTCGCTAAATCAGTCGAGTTAGATCGGTGACGGCGATTCTCACATTTGGAACGAGACC |
| 86 | GGTCTCA | CTGA | TTTATTCAGGGCACTACCCGGAGCT | AACGGGAGTCCGAGGTCGTACCTAG | ATCGGTGACGGCGATTCTCACATTT | GGAA | CGAGACC | 97 | 4 | 4_14 | GGTCTCACTGATTTATTCAGGGCACTACCCGGAGCTAACGGGAGTCCGAGGTCGTACCTAGATCGGTGACGGCGATTCTCACATTTGGAACGAGACC |
| 87 | GGTCTCA | CTGA | TTTATTCAGGGCACTACCCGGAGCT | GGATGGGAACCCAACACCTGTTCAT | ATCGGTGACGGCGATTCTCACATTT | GGAA | CGAGACC | 97 | 4 | 4_15 | GGTCTCACTGATTTATTCAGGGCACTACCCGGAGCTGGATGGGAACCCAACACCTGTTCATATCGGTGACGGCGATTCTCACATTTGGAACGAGACC |
| 88 | GGTCTCA | CTGA | TTTATTCAGGGCACTACCCGGAGCT | AGTGGTAGGTAGTTTTGCCCAGCCC | ATCGGTGACGGCGATTCTCACATTT | GGAA | CGAGACC | 97 | 4 | 4_16 | GGTCTCACTGATTTATTCAGGGCACTACCCGGAGCTAGTGGTAGGTAGTTTTGCCCAGCCCATCGGTGACGGCGATTCTCACATTTGGAACGAGACC |
| 89 | GGTCTCA | CTGA | TTTATTCAGGGCACTACCCGGAGCT | TTGGGCCGATAAGTTGAGTACCCTG | ATCGGTGACGGCGATTCTCACATTT | GGAA | CGAGACC | 97 | 4 | 4_17 | GGTCTCACTGATTTATTCAGGGCACTACCCGGAGCTTTGGGCCGATAAGTTGAGTACCCTGATCGGTGACGGCGATTCTCACATTTGGAACGAGACC |

| | | | | | | | | | | | |
|---|---|---|---|---|---|---|---|---|---|---|---|
| 90 | GGTCTCA | CTGA | TTTATTCAGGGCACTACCCGGAGCT | GCATAGGGTTCCACGCCAGTGTATG | ATCGGTGACGGCGATTCTCACATTT | GGAA | CGAGACC | 97 | 4 | 4_18 | GGTCTCACTGATTTATTCAGGGCACTACCCGGAGCTGCATAGGGTTCCACGCCAGTGTATGATCGGTGACGGCGATTCTCACATTTGGAACGAGACC |
| 91 | GGTCTCA | CTGA | TTTATTCAGGGCACTACCCGGAGCT | ATTCAGGGTTCCCACATCACGCAAA | ATCGGTGACGGCGATTCTCACATTT | GGAA | CGAGACC | 97 | 4 | 4_19 | GGTCTCACTGATTTATTCAGGGCACTACCCGGAGCTATTCAGGGTTCCCACATCACGCAAAATCGGTGACGGCGATTCTCACATTTGGAACGAGACC |
| 92 | GGTCTCA | CTGA | TTTATTCAGGGCACTACCCGGAGCT | GCATTCTGGTTCGCAGCTATATCGG | ATCGGTGACGGCGATTCTCACATTT | GGAA | CGAGACC | 97 | 4 | 4_20 | GGTCTCACTGATTTATTCAGGGCACTACCCGGAGCTGCATTCTGGTTCGCAGCTATATCGGATCGGTGACGGCGATTCTCACATTTGGAACGAGACC |
| 93 | GGTCTCA | CTGA | TTTATTCAGGGCACTACCCGGAGCT | GCATAACACAAGTACGGCTACGCAG | ATCGGTGACGGCGATTCTCACATTT | GGAA | CGAGACC | 97 | 4 | 4_21 | GGTCTCACTGATTTATTCAGGGCACTACCCGGAGCTGCATAACACAAGTACGGCTACGCAGATCGGTGACGGCGATTCTCACATTTGGAACGAGACC |
| 94 | GGTCTCA | CTGA | TTTATTCAGGGCACTACCCGGAGCT | GAAGGAGGCACATCCTTAAACCCGT | ATCGGTGACGGCGATTCTCACATTT | GGAA | CGAGACC | 97 | 4 | 4_22 | GGTCTCACTGATTTATTCAGGGCACTACCCGGAGCTGAAGGAGGCACATCCTTAAACCCGTATCGGTGACGGCGATTCTCACATTTGGAACGAGACC |
| 95 | GGTCTCA | CTGA | TTTATTCAGGGCACTACCCGGAGCT | GGCTATGTCCGTAACACTCCTAGCA | ATCGGTGACGGCGATTCTCACATTT | GGAA | CGAGACC | 97 | 4 | 4_23 | GGTCTCACTGATTTATTCAGGGCACTACCCGGAGCTGGCTATGTCCGTAACACTCCTAGCAATCGGTGACGGCGATTCTCACATTTGGAACGAGACC |
| 96 | GGTCTCA | CTGA | TTTATTCAGGGCACTACCCGGAGCT | GAGCGACAATCGACTTTCGTGGATC | ATCGGTGACGGCGATTCTCACATTT | GGAA | CGAGACC | 97 | 4 | 4_24 | GGTCTCACTGATTTATTCAGGGCACTACCCGGAGCTGAGCGACAATCGACTTTCGTGGATCATCGGTGACGGCGATTCTCACATTTGGAACGAGACC |

**Supplementary Table 4.** Primers used for DMX

| | Sequence (5' -> 3') |
|---|---|
| Forward library primer | atactacCGTCTCgaggaGGTGGATCAGGAGGTTCG |
| Reverse library primer | gcattacCGTCTCcggaaCCACTTCCACCGCTTCC |
| DMX 1_rv | TCATGCCGCGACGCATACTTTAGAC |
| DMX 2_fw | CTGGTTGGGATCAGTCGCTTAGTGC |
| DMX 3_rv | CTGGGTACTGTCCCAAGGACCACTC |
| DMX 4_fw | CGACACCGAACGTGCGACAAAACTA |
| DMX 5_rv | CTCTGCGACCTAGCCGGTTTATCCA |
| DMX 6_fw | TTTATTCAGGGCACTACCCGGAGCT |

**Supplementary Table 5.** Cost breakdown for DMX (green), and comparison to SAPP (blue) for the same number of designs.

| | **Library complexity** | | | |
|---|---|---|---|---|
| **Synthetic DNA** | **100** | **500** | **1000** | **2000** |
| Twist oligo pool (250-300 nt, list prices) | $ 1,030.00 | $ 2,060.00 | $ 3,090.00 | $ 4,121.00 |
| subtotal: | $ 1,030.00 | $ 2,060.00 | $ 3,090.00 | $ 4,121.00 |
| **Cloning** | | | | |
| BsaI | $ 76.26 | $ 152.52 | $ 305.05 | $ 610.10 |
| Salt T4 ligase | $ 115.20 | $ 230.40 | $ 460.80 | $ 921.60 |
| subtotal: | 191.46 | $ 382.92 | $ 765.85 | $ 1,531.70 |
| **Sequencing** | | | | |
| Minion flow cell (ONT) | $ 600.00 | $ 600.00 | $ 600.00 | $ 600.00 |
| Library Prep kit (ONT) | $ 95.00 | $ 95.00 | $ 95.00 | $ 95.00 |
| subtotal: | 695.00 | $ 695.00 | $ 695.00 | $ 695.00 |
| **Plasticware** | | | | |
| ECHO plates | $ 15.04 | $ 30.07 | $ 60.14 | $ 120.29 |
| 1536w rxn plates | $ - | $ 50.93 | $ 101.87 | $ 203.73 |
| 245mm Square BioAssay Dishes | $ 20.31 | $ 20.31 | $ 40.63 | $ 81.25 |
| subtotal: | 35.35 | $ 101.32 | $ 202.64 | $ 405.27 |
| **DMX Total Cost for the library ($)** | **1,951.81** | **3,239.24** | **4,753.49** | **6,752.97** |
| **DMX Cost / design ($)** | **19.52** | **6.48** | **4.75** | **3.38** |
| eBlocks Gene Fragments subtotal: | $ 2,100 | $ 10,500 | $ 21,000 | $ 42,000 |
| eBlocks Cloning subtotal: | $ 124 | $ 622 | $ 1,244 | $ 2,488 |
| **SAPP DNA Total cost ($)** | **2,224.40** | **11,122.00** | **22,244.00** | **44,488.00** |
| **SAPP DNA cost ($) / design** | **22.24** | | | |
| **Saving factor, DMX compared to eBlocks** | **1.14** | **3.43** | **4.68** | **6.59** |

**Supplementary Table 6.** Reagents and materials for SAPP.

| **Cloning** | **Supplier** | **Cat#** |
|---|---|---|
| BsaI-HF®v2 | NEB | R3733L |
| T4 DNA Ligase | NEB | M0202L |
| Entry vector (e.g. LM0627, Midiprep) | ZymoResearch | D4201 |
| BL21(DE3) Competent E. Coli | NEB | C2527I |
| NEB Stable for GGA vector propagation with *ccdb* lethal gene | NEB | C3040H |
| **Protein Expression** | | |
| Terrific broth II | MP Biomedicals | 113046032 |
| Glycerol | MilliporeSigma | G5516-4L |
| Dextrose (D-Glucose), Anhydrous | FisherScientific | D16-1 |
| D-Lactose monohydrate | MilliporeSigma | 61345-1KG |
| Kanamycin sulfate | FisherScientific | AC450811000 |
| **Protein Purification** | | |
| B-PER™ | ThermoFisher | 78248 |
| Benzonase® Nuclease | MilliporeSigma | E1014-25KU |
| PMSF | MilliporeSigma | 11359061001 |
| Lysozyme from chicken egg white | MilliporeSigma | L6876 |
| HisPur™ Ni-NTA Resin | ThermoFisher | 88223 |
| Cytiva S200/S75 Increase 5-150 | Cytiva | 29148722 |
| **Buffers** | | |
| Trizma® base | MilliporeSigma | T1503-10KG |
| NaCl | FisherScientific | BP358-212 |
| Imidazole | MilliporeSigma | 1370981000 |
| **Plasticware** | | |
| Tips | Opentrons | 999-00013 |

| | | |
|---|---|---|
| Armadillo PCR plate | FisherScientific | AB2396 |
| 96-well 2 mL Deepwell culture plate | FisherScientific | 278752 |
| Breathable membrane (Diversified Biotech Breathe Easier) **Do not use the "Breathe Easy" from the same manufacturer, we had poor expression results with the "Breath Easy" film.** | MilliporeSigma | Z763624 |
| 96-well fritted plate 25 µm | Agilent | 200967-100 |
| 96-well filter plate 0.2 µm | Agilent | 203940-100 |
| 96-well receiver plate | ThermoFisher | 249944 |
| 384-well HPLC collection plate | Greiner Bio-One | 781270 |

**Supplementary Table 7**. SAPP Buffers.

| Name | Recipe | amount | comment |
|---|---|---|---|
| **TB-2 Media** | from MP Biomedicals | 50 g/L | prepare in 1L flasks, autoclave |
| **5052 (50x stock)** | Glycerol | 250 g/L | sterile filter |
| | D-(+)-Glucose monohydrate | 27.5 g/L | |
| | α-Lactose monohydrate | 100 g/L | |
| **Autoinduction media (1 L)** | 50x 5052, sterile | 20 mL | prepare sterile (under flame) |
| | 1 M $MgSO_4$, sterile | 2 mL | |
| | 1000 x antibiotic, sterile | 1 mL | e.g. Kanamycin final 50 µg/mL |
| | TB2 | to 1 L | |
| **HIS WASH BUFFER** | Tris | 20 mM | pH 8 |
| | NaCl | 300 mM | |
| | Imidazole | 25 mM | |
| **HIS ELUTION BUFFER** | Tris | 20 mM | pH 8 |
| | NaCl | 300 mM | |
| | Imidazole | 500 mM | |
| **His resin stripping buffer** | NaOH | 500 mM | |
| | EDTA-2Na·$2H_2O$ | 50 mM | |
| **SNAC cleavage buffer** | CHES | 100 mM | pH 8.6 |
| | Acetone Oxime | 100 mM | |
| | NaCl | 100 mM | |
| | Guanidinium HCl | 500 mM | |

**Supplementary Table 8.** Reagents and materials for DMX.

| **Cloning** | **Supplier** | **Cat#** |
|---|---|---|
| KAPA HiFi HotStart ReadyMix | Roche | KK2602 |
| EvaGreen Dye | Biotium | 31000 |
| Zymoclean Gel DNA Recovery Kits | Zymo Research | D4007/D4008 |
| BsmBI-v2 NEBridge Golden Gate Assembly kit | NEB | E1602S |
| DNA Clean & Concentrator-5 | Zymo Research | D4003 |
| E. Cloni EXPRESS BL21(DE3) electrocompetent cells | Lucigen | 60300-1 |
| BsaI-HF®v2 | NEB | R3733L |
| Salt-T4 DNA Ligase | NEB | M0467L |
| Zyppy Plasmid Miniprep | Zymo Research | D4036 |
| dsDNA Quantitation, High Sensitivity | Invitrogen | Q32851 |
| **Plasticware** | | |
| Breathe Easier Plate Seal | Fisher Scientific | NC1664397 |
| Axygen® Foil Plate Seal | Fisher Scientific | PCR-AS-600 |
| ECHO-qualified 384-well plates | Beckman Coulter | C74290 |
| ECHO-qualified polypropylene 1536-well plate | Greiner | 782270 |
| Bioassay plates | Corning | 431111 |

**Supplementary Table 9.** CryoEM data collection, refinement, and validation statistics

| | RSV-F SC-DM/cb13 PDB:9YCF EMDB-72769 |
|---|---|
| Data collection and processing | |
| Magnification | 36,000 × |
| Voltage (kV) | 200 |
| Electron exposure (e–/Å$^2$) | 50 |
| Defocus range (μm) | 1-2 |
| Pixel size (Å) | 0.4425 |
| Symmetry imposed | C1 |
| Initial particle images (no.) | 866,428 |
| Final particle images (no.) | 89,296 |
| Map resolution (Å) | 4.62 |
| FSC threshold | 0.143 |
| Map resolution range (Å) | 4.0-6.0 |
| Refinement | |
| Initial model used (PDB code) | |
| Model resolution (Å) | 4.62 |
| FSC threshold | 0.143 |
| Model resolution range (Å) | 4.0-6.0 |
| Map sharpening *B* factor (Å$^2$) | 217.3 |
| Model composition | |
| Non-hydrogen atoms | 7045 |
| Protein residues | 1423 |
| Ligands | N/A |
| *B* factors (Å$^2$) | |
| Protein | 178.59 |
| Ligand | N/A |
| R.m.s. deviations | |
| Bond lengths (Å) | 0.002 |
| Bond angles (°) | 0.525 |
| Validation | |
| MolProbity score | 1.39 |
| Clashscore | 2.18 |
| Poor rotamers (%) | 0.0 |
| Ramachandran plot | |
| Favored (%) | 94.11 |
| Allowed (%) | 5.89 |
| Disallowed (%) | 0 |
